# SIX1 Reprograms Myogenic Transcription Factors to Maintain the Rhabdomyosarcoma undifferentiated state

**DOI:** 10.1101/2021.04.25.439216

**Authors:** Jessica Y. Hsu, Etienne P. Danis, Stephanie Nance, Jenean O’Brien, Veronica M. Wessells, Andrew E. Goodspeed, Jared C. Talbot, Sharon L. Amacher, Paul Jedlicka, Joshua C. Black, James C. Costello, Adam D. Durbin, Kristin B. Artinger, Heide L. Ford

**Affiliations:** Department of Pharmacology, University of Colorado Anschutz Medical Campus, Aurora, CO, USA; Pharmacology Graduate Program, University of Colorado Anschutz Medical Campus, Aurora, CO, USA; Department of Biology, College of St. Scholastica, Duluth, MN, USA; Division of Medical Oncology, University of Colorado Anschutz Medical Campus, Aurora, CO, USA; School of Biology and Ecology, University of Maine, Orono, ME, USA; Department of Molecular Genetics, Ohio State University, Columbus, OH, USA; Department of Pathology, University of Colorado Anschutz Medical Campus, Aurora, CO, USA; Division of Molecular Oncology, St. Jude Children’s Research Hospital, Memphis, TN, USA; Department of Craniofacial Biology, University of Colorado Anschutz Medical Campus, Aurora, CO, USA; University of Colorado Cancer Center, University of Colorado Anschutz Medical Campus, Aurora, CO, USA

**Keywords:** Rhabdomyosarcoma, SIX1, MYOD1, muscle differentiation, muscle progenitor, transcriptional control

## Abstract

Rhabdomyosarcoma (RMS) is a pediatric skeletal muscle sarcoma characterized by the expression of the myogenic-lineage transcription factors (TF) MYOD1 and MYOG. Despite high expression of these TFs, RMS cells fail to terminally differentiate, suggesting the presence of factors that alter their function. Here, we demonstrate that the developmental TF, SIX1, is highly expressed in RMS and is critical to maintain a muscle progenitor-like state. SIX1 loss induces terminal differentiation of RMS cells into myotube-like cells and dramatically impedes tumor growth *in vivo*. We show that SIX1 maintains the RMS undifferentiated state by controlling enhancer activity and MYOD1 occupancy at loci more permissive to tumor growth over terminal muscle differentiation. Finally, we demonstrate that a gene signature derived from SIX1 loss correlates with differentiation status in RMS and predicts RMS progression in human disease. Our findings demonstrate a master regulatory role for SIX1 in the repression of RMS differentiation via genome-wide alterations in MYOD1-mediated transcription.

**Highlights:**

- SIX1 prevents differentiation in RMS while it promotes differentiation during normal development
- FN-RMS are highly dependent on SIX1 for growth in both zebrafish and mouse xenograft models
- Loss of SIX1 alters the transcriptional landscape of RMS cells, inducing a growth to differentiation switch
- SIX1 knockdown in FN-RMS causes reduced super enhancer-based activity at stem-related genes and enhanced MYOD1 binding to differentiation loci, resulting in the activation of a myogenic differentiation program
- A gene signature derived from SIX1 loss strongly correlates with myogenic differentiation status and is predictive of advanced RMS.

## Introduction

Rhabdomyosarcoma (RMS) is a soft tissue pediatric sarcoma with molecular and histological features that resemble undifferentiated skeletal muscle. The majority of pediatric RMS cases can be divided into two major subtypes: Embryonal RMS (ERMS) and Alveolar RMS (ARMS), which are designated based on their histology. While ERMS tumors are characterized by a variety of mutational events, notably *RAS* mutations, ARMS tumors are classically associated with PAX3-FOXO1 or PAX7-FOXO1 chromosomal rearrangements, which has led to the replacement of the histological annotations ERMS and ARMS with “Fusion-negative (FN)” and “Fusion-positive (FP)”. The distinct genetic perturbations associated with ERMS and ARMS have long implied that the RMS subtypes arise from distinct mechanisms, however a shared feature of all RMS tumors is their expression of the myogenic regulatory transcription factors (TF) *MYOD1* and *MYOG*, orchestrators of skeletal muscle differentiation with aberrant functions in RMS tumors^1^. Whereas in normal skeletal muscle differentiation these myogenic TFs coordinate the expansion, commitment, and eventual differentiation of embryonic mesodermal or myogenic progenitors, the expression of these myogenic TFs in RMS tumors is not coupled with exit from the cell cycle and differentiation into post-mitotic myofibers^2^. Several studies to date have discovered distinct activities of these myogenic transcription factors in the context of normal muscle development and RMS^3–5^. However, it remains less clear what factors cause these myogenic regulatory factors to depart from their canonical roles as drivers of muscle differentiation to instead maintain RMS cells as less differentiated muscle progenitors.

The *SIX1* homeodomain-containing TF belongs to the *Six* gene family that includes *SIX1-SIX6* in vertebrates. Early studies of the *SIX1* ortholog in drosophila, *sine oculis (so)*, placed the functions of the *Six* gene family in eye morphogenesis, as *so* mutants lack compound eye structures^6^. However, since the original discovery of *so,* the functions of the *Six* family genes are known to extend beyond the visual system in vertebrates. Notably, the mammalian orthologs *Six1* and *Six4* have conserved and indispensable roles in embryonic skeletal muscle development and skeletal muscle regeneration. In mice, *Six1* deficiency alone causes reduced and disorganized muscle mass^7^, and further ablation of *Six1* and its ortholog *Six4* causes exacerbated craniofacial defects and severe muscle hypoplasia^8^. In both *Six1* and *Six1/Six4* deficient mouse models, the expression of the critical myogenic TFs *MYOD1* and *MYOG* is compromised in migrating hypaxial muscle, demonstrating that *Six1* and *Six4* are required for the activation of these myogenic TFs. In zebrafish, morpholino-mediated loss of *six1b* gene expression similarly causes reduced hypaxial muscle and impairment of *Pax7+* muscle stem cell proliferation during skeletal muscle repair^9, 10^. Recently, genetic ablation of *six1a/six1b/six4a/six4b* paralogs in the zebrafish genome has additionally shown that compound loss of *six1/4* function causes complete loss of all migratory muscle precursors that generate hypaxial muscles such as the fin muscles, while leaving trunk muscle relatively unaffected^11^. These results align with previous observations that morpholino-mediated loss of *six1a* and *six1b* also affect hypaxial muscles, though the muscle defects observed in the morpholino studies are more severe than those seen in the *six1a/six1b* genetic mutant^9–11^. These studies demonstrate that *Six1*, which acts in concert with *Six4*, lies upstream of the myogenic specification gene regulatory network and is a necessary component of the skeletal muscle circuit.

Myogenic differentiation is tightly governed by a cascade of myogenic regulatory factor (MRF) expression which encompass the highly conserved class II basic helix-loop-helix (bHLH) TFs *MYOD1, MYF5, MYOG,* and *MRF4.* During the course of embryonic development as well as skeletal muscle repair and regeneration, these four MRFs are considered necessary for committing progenitor cells to the skeletal muscle lineage, expanding the progenitor cell pool, and differentiating committed cells into contractile muscle fibers^12^. While structurally the MRF family is conserved, the transition of muscle progenitors from commitment, to growth, and subsequently to differentiation invokes sub-functionalized and context- specific roles of these MRFs. Indeed, *MyoD1* can activate distinct myoblast-specific and differentiation- specific gene expression programs by modifying chromatin environments that facilitate either differentiation or myoblast growth^13, 14^. Because the functions of *MYOD1* are co-opted in RMS tumors to foster growth rather than to promote differentiation, we hypothesized that other factors critical for normal skeletal muscle development must repress the differentiation subprograms of *MYOD1.* Given the well- established role of *SIX1* in regulating upstream activities of *MYOD1* as well as other MRFs to induce skeletal muscle development^8, 10, 15–17^, we investigated the molecular role of *SIX1* in regulating RMS tumor growth. Here, we report that SIX1 loss causes a growth-to-differentiation switch in RMS cells by globally regulating a myogenic transcriptional program and reinstating the function of MYOD1 as a driver of skeletal muscle differentiation.

## Results

### SIX1 is overexpressed and predicted to be an essential gene in Rhabdomyosarcoma

To examine whether SIX1 is highly expressed in human RMS, we interrogated its expression in publicly available large RMS RNAseq datasets. In multiple independent datasets, high SIX1 mRNA expression could be detected, both compared with other sarcomas in the National Cancer Institute Oncogenomics pan-sarcoma dataset (Suppl Fig. 1A) and the St. Jude Pediatric Cancer Genome Project (Suppl Fig. 1B), and compared with normal tissues in the St. Jude Integrated Rhabdomyosarcoma Database (iRDb) (Fig 1A). Notably, SIX1 was more highly expressed in RMS samples, compared with differentiated skeletal muscle controls depicting different stages of skeletal muscle development (Fig 1A). To confirm these data, we next assessed SIX1 protein expression in an RMS tumor tissue array consisting of 96 human RMS patient samples and 8 normal skeletal muscle controls (Figure 1B-C). Using a 1-4 scoring system of nuclear immunohistochemical staining, we detected strong nuclear SIX1 staining in the ERMS/Fusion-Negative and ARMS/Fusion-Positive tumor sections (18% and 29% with IHC staining scores ≥ 2, respectively) compared to normal skeletal muscle control sections (0% with IHC staining score ≥2) (Figure 1B-C). To further determine if SIX1 has a functional role in RMS, we next examined data from the Broad and Sanger Institutes’ exome-wide CRISPR-Cas9 knockout (KO) screening dataset^18^. In the 869 cell lines tested in the CRISPR-Cas9 screen, we observed that the 10 RMS cell lines used in the screen exhibited both high SIX1 mRNA expression and high SIX1 gene dependency (Figure 1D). Further comparison of the RMS tumor cell lines against all other tumor cell lines demonstrates that SIX1 is a selective dependency in RMS and is required for RMS cell survival (*q-*value = 0.018), as is the myogenic TF MYOD1 (Figure 1E).

**Figure 1.**
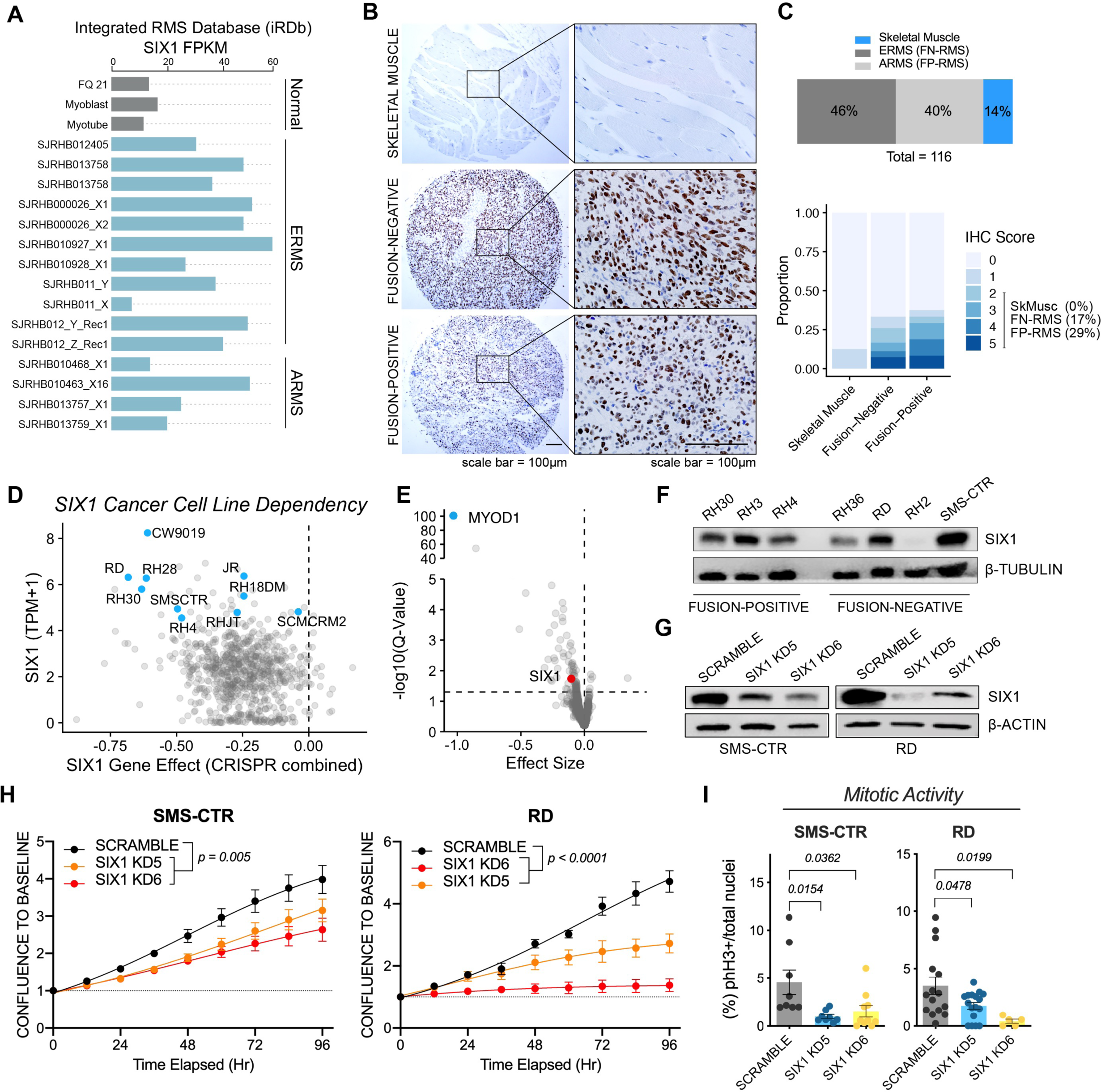
SIX1 is overexpressed and predicted to be an essential gene in Rhabdomyosarcoma. (A)) Fragments per kilobase million (FPKM) expression of SIX1 in the St. Jude Pediatric Cancer Genome Project cohort (Grey = three normal skeletal muscle controls; FQ21 = fetal quadricep muscle). (B) IHC staining counterstained with hematoxylin and DAB intensity scoring of an RMS tumor array with normal skeletal muscle controls. (C) Frequency distribution of IHC scores across RMS and skeletal muscle tissue cores and frequency distribution of tissue cores with IHC scores ≥2. (D) SIX1 transcripts per kilobase million (TPM) expression against SIX1 gene effect score in 1775 cell lines in the Cancer Dependency map CRISPR-Cas9 large-scale KO screen (RMS cell lines in blue). (E) Volcano plot of gene dependency scores for MYOD1 (blue) and SIX1 (red) in RMS cell lines versus all other cell lines of different tissue types. Statistical analysis of gene dependencies between RMS and all other cell types were performed using a two-class Kolmogorov-Smirnov test. (F) Western blot of SIX1 protein levels across a panel of FN and FP RMS human cell lines (G) shRNA-mediated knockdowns of SIX1 in RD and SMS-CTR cell lines. (H) IncuCyte live-cell imaging growth assays of SMS-CTR and RD shScramble and SIX1 KD cells over a 96-hrs. Cells were plated in triplicate and relative cell growth was measured by normalizing cell confluency at each time point relative to initial timepoint confluency. Data represent mean ± SEM and statistical differences between shScramble and SIX1 KD5 or SIX1 KD6 was measured by fitting data to a longitudinal mixed effects model. (I) Mitotic activity of SMS-CTR and RD shScramble and SIX1 KD cells measured by phH3 staining. Cells were counterstained with DAPI. Data represent mean ± SEM of at least 3 independent experiments.

Given the high expression of SIX1 in RMS tumors compared with matched normal tissues, we hypothesized that the increased SIX1 expression in RMS tumors compared with normal muscle could aberrantly activate its developmental functions in this cancer context. To investigate SIX1 function in RMS, we examined the expression of SIX1 in a panel of human RMS cell lines and detected high SIX1 expression in both FN and FP RMS cell lines (Figure 1F). Although SIX1 expression is high in both FP and FN-RMS, we focused our studies on the FN subtype to interrogate its functions outside the context of the *PAX3-FOXO1* fusion. Using two FN-RMS cell lines (SMS-CTR and RD) that highly express SIX1, we sought to validate the CRISPR-Cas9 screen findings using an orthogonal method. We thus established SMS-CTR and RD cell lines transduced with shRNAs targeting either no coding sequence in the genome (shScramble) or two distinct SIX1 sequences (SIX1 KD5, SIX1 KD6) (Figure 1G). In both cell lines, we observed that reduced levels of SIX1 were paired with deficits in cell growth and mitotic activity as measured by IncuCyte live-cell growth assays (Figure 1H) and the mitotic marker phospho-histone H3 (phH3) staining, respectively (Figure 1I). Together, these data demonstrate that SIX1 is highly expressed and required for the growth of RMS cells *in vitro*.

### *six1b* is required for zebrafish RMS tumor growth

Given the above *in vitro* observations, we sought to examine the role of SIX1 in an *in vivo* setting, first using a zebrafish model of ERMS (zRMS) induced by the co-injection of *rag2-kRASG12D* and *rag2-GFP* transgenes into the single-cell stage of the zebrafish ^19^. This model results in the generation of skeletal muscle tumors with histological features similar to human FN-RMS, and parallels our cell line data, as SMS-CTR and RD cells are both *RAS*-mutated FN-RMS^20, 21^. To examine the expression of the two zebrafish *six1* paralogs, *six1a* and *six1b,* in zRMS tumors, we performed quantitative real-time PCR (qRT-PCR) analysis and found that *six1b* was significantly upregulated in zRMS tissue compared to age-matched normal skeletal muscle (Figure 2A), which was confirmed using RNA *in-situ* hybridization (ISH) (Figure 2B). To determine whether *six1b* was required for RMS tumor growth *in vivo,* we then combined the zRMS injection model with zebrafish carrying genetic loss-of-function alleles for only *six1b*^11^, both because of its more consistent overexpression in zRMS, and because the *six1a;six1b* double mutant fails to survive to adult stages when zRMS tumors would typically form^11, 19^. In contrast, *six1b* mutants develop normally and are therefore a suitable model to test the function of reduced *six1* levels in RMS *in vivo.* Consistent with our previous findings, we found no differences in *pax3a, myod1* or *myogenin* expression between wildtype and *six1b* mutant sibling embryos from the 5-20+ somite stages (Suppl Fig 2A-C)^11^.

**Figure 2.**
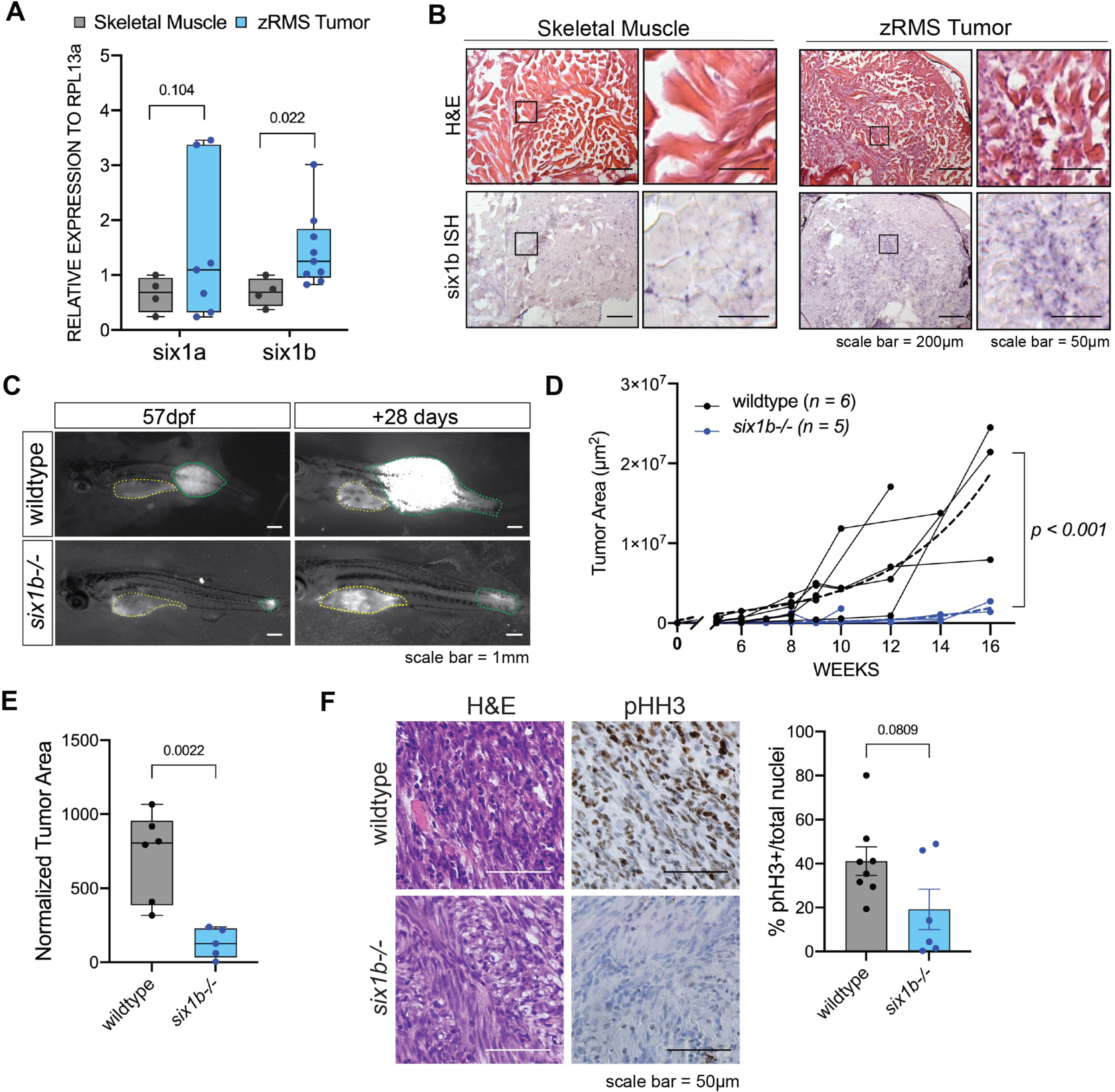
*six1b* is required for zebrafish RMS tumor growth. (A) qRT-PCR expression of zebrafish *six1* paralogs *six1a* and *six1b* in dissected GFP+ zRMS tumor tissue compared to age-matched normal skeletal muscle (*n* = 4 normal muscle samples, *n* = 6 zRMS tumor samples). (B) Representative images of *six1b* transcripts as visualized by H&E and *in situ* hybridization signal (purple puncta; *n = 5* fish per group) (C) Representative images of tumor progression (outlined in green) over 28 days from 57-85 days post fertilization (dpf) between wildtype and *six1b^-/-^* tumor-burdened individuals. Yellow outline represents autofluorescence from stomach and yolk. (D) Quantification of tumor area by GFP+ tumor area in each individual fish over time. Tumor growth per individual is represented as individual tracks and composite growth of wildtype and *six1b^-/-^* tumors was fitted to a non-linear logistical growth model and represented by dotted lines. A longitudinal mixed effect model was used to measure statistical differences between conditions over repeated measures. (E) Tumor area growth over time normalized to standard length of fish at 120 dpf or at prior time point due to moribundity. (F) Representative staining and quantification of H&E and phospho- histone H3 IHC (brown) in sectioned zRMS tumors. Dots in graph represent %phH3 staining per tumor section; phH3 staining quantified over 2 sections per tumor (*n =* 4 wt tumors, *n =* 3 *six1b^-/-^* tumors). Statistical differences were calculated using a Welch’s *t-*test.

To determine whether *six1b* loss is sufficient to alter *kRAS-*mediated zRMS tumorigenesis, we injected *rag2-kRASG12D/GFP* transgenes^19^ into the progeny of *six1b*^+/-^ breeding pairs to generate age-matched sibling groups with all possible *six1b* genotypes. Interestingly, while GFP positivity could be detected in all genotypes, the progression to overt tumors was largely lost with *six1b* depletion (Figure 2C-E). Following tumor initiation, however, we observed that tumors established in *six1b^-/-^* zebrafish grew significantly slower over a 120-day time course, as compared to tumors established in wildtype siblings (Figure 2C-D). Reflecting this reduced growth rate, *six1b^-/-^* tumors were smaller in size compared to that of wildtype siblings’ tumors at their final collection time-point at 120 dpf (Figure 2E). Immunohistochemical staining of tumors demonstrated that while wildtype tumors displayed normal architecture of RMS, *six1b^- /-^* tumor cells displayed more elongated morphology with higher cytoplasmic to nuclear ratios, reminiscent of skeletal muscle differentiation (Figure 2F). In alignment with the slow growth rate, staining for phH3 in *six1b^-/-^* (*n* = 3) tumors trended toward lowered intensity when compared to prominent phH3-positive staining in wildtype zRMS tumors (*n* = 4). This downward shift did not reach statistical significance (*p=0.081*) likely due to the to the small number of *six1b^-/-^* tumors that formed and were evaluable.

Nevertheless, the reduction in GFP+ tumor growth in the *six1b^-/-^* zebrafish indicate that *six1b* plays a critical role in zRMS tumor progression at least in part via controlling RMS tumor cell proliferation.

### SIX1 knockdown inhibits human RMS tumor growth and progression

The tumors that formed in *six1b^-/-^* zebrafish displayed an elongated, more spindle-cell morphology, suggesting that RMS cell-state fundamentally differs between RMS cells derived from wildtype and *six1b* depleted animals. To identify whether similar changes occur in human RMS, we examined the morphology of SMS-CTR and RD cells that were transduced with SIX1 shRNAs. Within approximately five passages after stable SIX1 KD, both RMS cell lines began to exhibit an altered, elongated morphology, distinguishing them from shScramble controls cells (Figure 3A-B).

**Figure 3.**
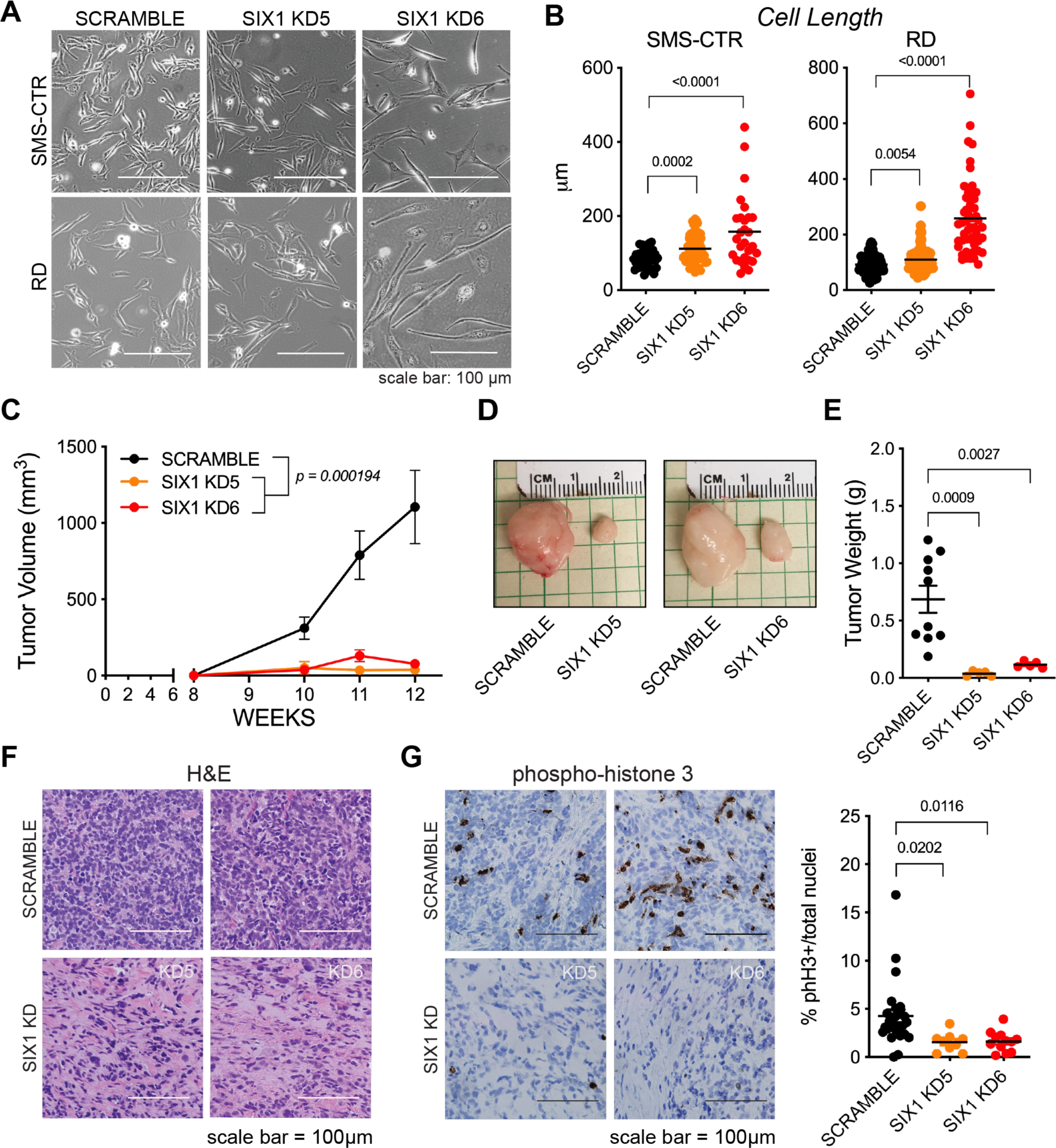
SIX1 knockdown inhibits human RMS tumor growth and progression. (A) Brightfield images depicting elongated cell morphology of SIX1 KD SMS-CTR and RD cells along with (B) quantification of cell lengths. (C) Tumor volumes, measured by caliper, over a 12-week time period of shScramble and SIX1 KD SMS-CTR cells that were engrafted bilaterally into the flanks of NOD/SCIDγ (NSG) mice. Data represent mean ± SEM and were fitted to a longitudinal mixed effects model for statistical analysis of shScramble and SIX1 KD samples (D) Representative images of dissected shScramble or SIX1 KD xenografted tumors at 12 weeks. (E) Final tumor weights in grams at the end of the 12-week study. (*n* = 10 mice total; 10 mice received shScramble cells in one flank, and SIX1 KD cells in opposite flank. 5 mice received a SIX1 KD5 flank injection, and 5 mice received a SIX1 KD6 flank injection). (F) Representative H&E histology of dissected shScramble and SIX1 KD xenografted tumors. (G) Representative phH3 immunostaining (brown) of dissected shScramble and SIX1 KD xenografted tumors. Dots in graph represent %phH3+ staining per tumor section; phH3 staining quantified over 2 sections per tumor.

We next assessed the *in vivo* outcomes of SIX1 KD in RMS tumor growth. SMS-CTR shScramble and SIX1 KD cells were xenografted subcutaneously in Matrigel into either the left or right flank of immune-compromised *NOD/SCID/IL2Rγ* mice and screened weekly for tumor growth. Tumor growth over time, as represented by tumor volume and final tumor weight, was significantly reduced in SIX1 KD tumors compared to shScramble tumors (Figure 3C-E). Histological characterization of the dissected control and SIX1 KD tumors by H&E revealed clear histological distinctions between shScramble and SIX1 KD tumors whereby all shScramble tumors exhibited high cell density while SIX1 KD tumors were sparsely populated with cells distinguished by elongated nuclear and cytoplasmic morphology (Figure 3F). Notably, upon staining xenografted tumors for phH3, we found that SIX1 KD tumors exhibited significantly less mitotic activity than shScramble tumors (Figure 3G), yet apoptosis, as measured by cleaved caspase 3 (CC3) staining, was unchanged (Suppl Fig 3). These data demonstrate that the profound differences in *in vivo* tumor growth between shScramble and SIX1 KD RMS tumors can be largely attributed to the lower proliferative capacity of SIX1 KD tumors, and are not due to higher levels of apoptosis.

### SIX1 knockdown induces myogenic differentiation in RMS cells

As described above, loss of SIX1 suppresses *in vitro* and *in vivo* RMS growth, and leads to alterations in cell morphology, consistent with morphological changes that occur during myogenic differentiation. Because SIX1 KD induced profound cell elongation and anti-proliferative phenotypes in our RMS cell lines, we asked whether these phenotypes were a consequence of SIX1 directly regulating a pro- proliferative transcriptional program, or a secondary consequence of another upstream program regulated by SIX1. We hypothesized that similar to its functions in normal skeletal muscle development, SIX1 overexpression in RMS may regulate an early myogenic transcriptional program that supports RMS cell proliferation and self-renewal^7, 17^. Therefore, to delineate the transcriptional program coordinated by SIX1 in RMS, we performed RNA-sequencing analysis (RNAseq) on our SMS-CTR shScramble and SIX1 KD cell lines.

The RNAseq analysis revealed a total of 1017 differentially expressed genes (|Fold-change| ≥ *1.5 & FDR ≤ 0.25*) between SMS-CTR shScramble and SIX1 KD cells (Figure 4A). Of note, numerous muscle-specification genes such MYOG, MYMK, and MYMX, were marked as significantly upregulated while genes known to regulate cell motility and invasion such as TWIST2 and L1CAM were significantly downregulated^22–24^. To identify dysregulated pathways upon SIX1 KD, we performed gene set enrichment analysis (GSEA)^25^. This analysis revealed an overarching positive enrichment of muscle cell differentiation and contractile muscle gene signatures in SIX1 KD cells (Figure 4B) while chromatin assembly and developmental cell growth signatures were negatively enriched in SIX1 KD cells (Figure 4B, Suppl Fig 4A). Upon closer inspection of gene expression within the MSigDB Myogenesis hallmark pathway, we again observed a clear switch in the expression pattern of canonical myogenic genes from low expression in shScramble cells to higher expression in SIX1 KD cells (Figure 4C).

**Figure 4.**
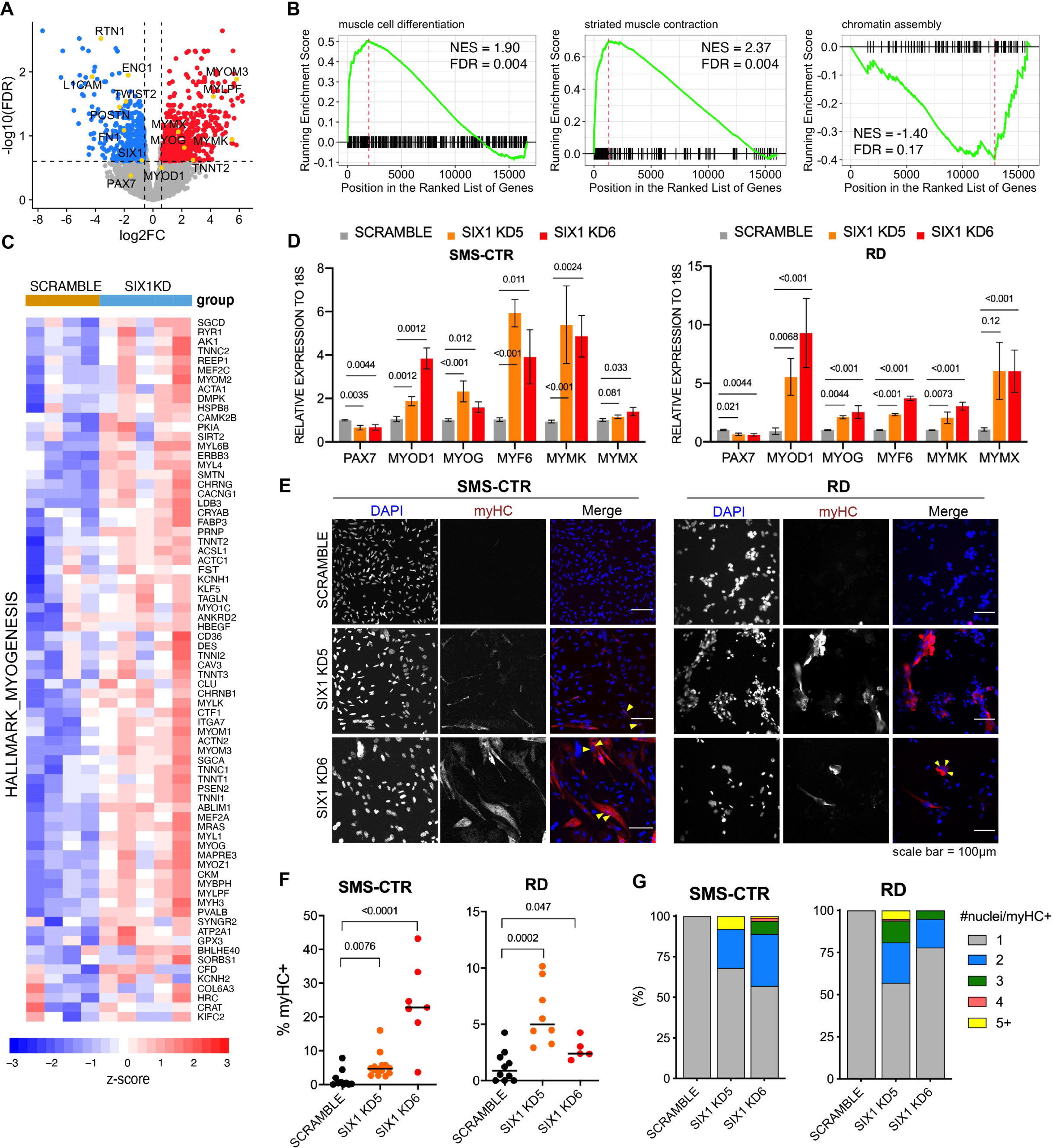
SIX1 knockdown induces myogenic differentiation in RMS cells. (A) Volcano plot of log2fold-change (FC) gene expression (SIX1 KD over shScramble) and adjusted *p-*value after edgeR-based differential expression analysis from the SMS-CTR RNAseq experiment. Red dots denote genes significantly upregulated (FC≥1.5 & adj p-value ≤0.25) and blue dots denote genes significantly downregulated (FC≤-1.5 & adj p-value ≤0.25) upon SIX1 KD. (B) Gene set enrichment analysis plots of ranked log2FC expression (SIX1 KD over shScramble) show positive enrichment for curated muscle cell differentiation and skeletal muscle contraction gene signatures and negative enrichment for chromatin assembly gene signatures. (C) Heatmap plotting expression of the MSigDB myogenesis gene set across shScramble and SIX1 KD samples. Scale bar represents z-score- converted log2CPM values. (D) Validation of differential mRNA expression of genes involved in muscle differentiation in SMS-CTR and RD cell lines with SIX1 KD by qRT-PCR. Barplot data represent mean ± SEM expression values across *n* ≥ 5 independently collected biological samples. (E) Positive MyHC (MF-20, red) immunostaining and DAPI counterstain (blue) in SIX1 KD RMS cells compared to shScramble RMS cells. (F) Quantification of myHC staining over total nuclei per field of view (each dot represents %myHC+ cells over one technical replicate from at least 3 independent experiments) and (G) fusion indices of SMS-CTR and RD control and SIX1 KD cells.

To validate the changes observed in SIX1 KD cells by RNAseq, we performed qRT-PCR in both SMS-CTR and RD cell lines for a subset of differentially expressed myogenic genes identified from our RNAseq analysis. Compared to their respective shScramble control cells, SMS-CTR and RD SIX1 KD cells expressed reduced levels of *PAX7* (a TF enriched in muscle progenitors) and expressed higher levels of the myogenic regulatory factors *MYOD1, MYOG,* and *MYF6.* In agreement with our RNAseq results, we also observed increased expression of genes associated with myoblast fusion: *MYMK* and *MYMX* (Figure 4D)^28^. To further examine whether our SIX1 KD cells underwent myogenic differentiation, we stained SMS-CTR and RD SIX1 KD cells for myosin heavy chain (myHC), a marker of terminal muscle differentiation. In both cell line models, SIX1 KD cells exhibited higher proportions of myHC+ cells (Figure 4E-F) and were more frequently multinucleated than shScramble cells (Figure 4G). These data indicate that SIX1 KD RMS cells are capable of terminally differentiation and forming multinucleated myofibers in contrast to shScramble cells, which maintain their muscle progenitor state.

To determine whether this muscle differentiation phenotype observed with SIX1 loss in human RMS models is conserved in the zRMS model, we additionally stained wildtype and *six1b^-/-^* zRMS tumors for Pax7 and myHC. In evaluable wildtype and *six1b^-/-^* tumor sections, we observed a decrease in Pax7 staining in *six1b^-/-^* tumors compared to wildtype tumors (Suppl Fig 5A), indicative of a shift in differentiation status of the tumors toward a more myotube-like state. In one particular *six1b^-/-^* tumor, we observed strong myHC staining in the tumor section which contrasted the largely absent myHC staining in all wildtype tumor sections (Suppl Fig 5B). Taken together, these data demonstrate that *SIX1* functions to repress a myogenic differentiation program in RMS cells in both human and zebrafish models.

**Figure 5.**
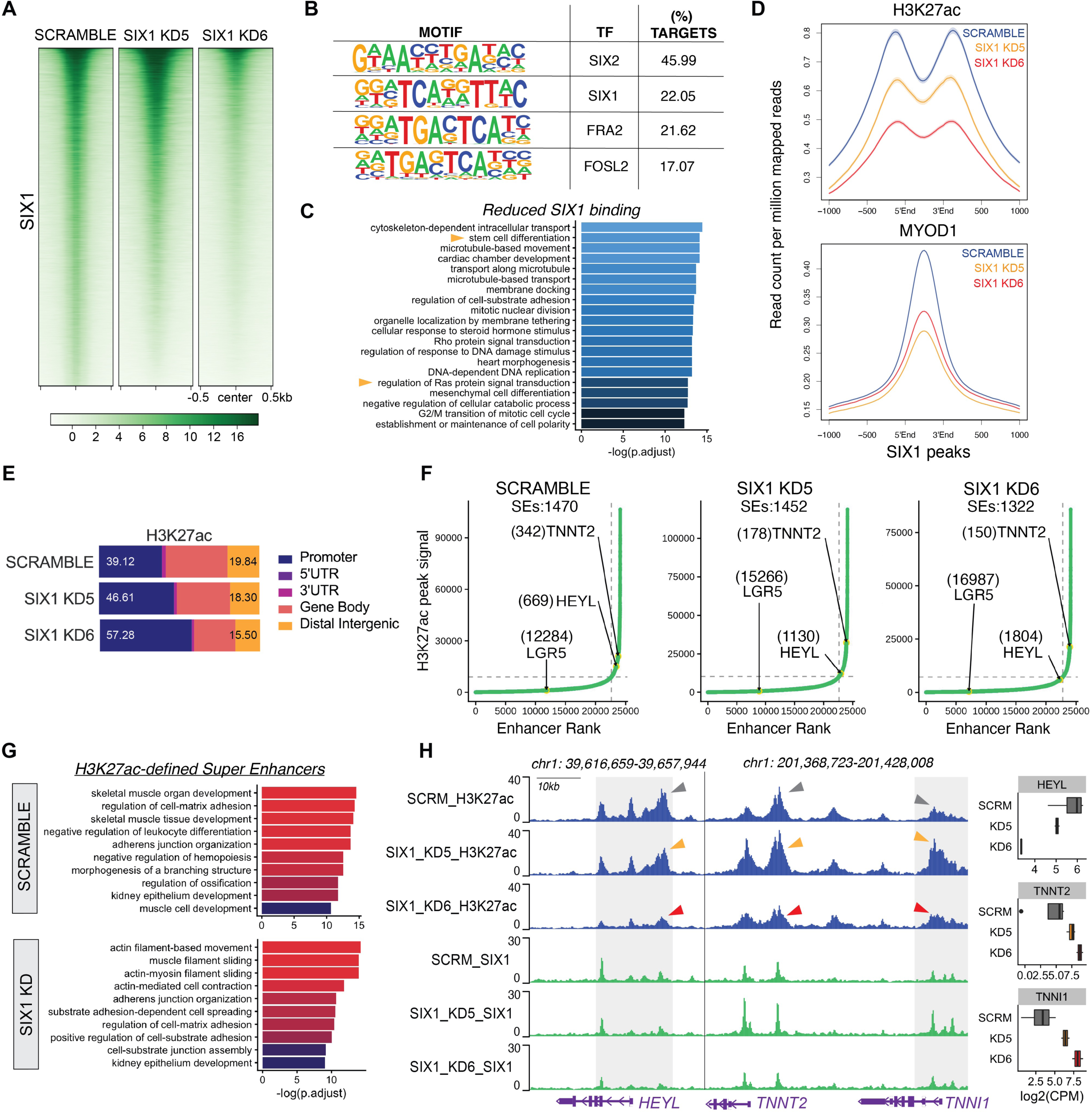
SIX1 globally regulates both stem/oncogenic and myogenic differentiation genes through fine-tuning of super-enhancer activity. (A) Heatmaps of genome-wide SIX1 ChIPseq signal in SMS-CTR shScramble, SIX1 KD5, and SIX1 KD6 cells. Heatmaps were generated using deepTools and centered at shScramble SIX1 peaks. (B) Motif analysis on peak coordinates exhibiting reduced 1.5-fold SIX1 binding in both SIX1 KD5 and SIX1 KD6 SMS-CTR SIX1 ChIPseq datasets. Top 4 enriched motifs shown. (C) Pathway enrichment plots of annotated sites of SIX1 loss in both SIX1 KD5 and KD6 cell lines. Enrichment plots were generated using ChIPseeker followed by ClusterProfiler R packages with gene set sizes restricted to 100 to 250 genes and a q-value cut-off of 0.05. (D) ChIPseq average profiles of MYOD1, and H3K27ac signal over SIX1 binding sites that exhibited reduced binding in SIX1 KD cells compared to shScramble cells. Average profiles were centered around reduced SIX1 peaks and show co-occurrence of SIX1 and MYOD1 binding as well as H3K27ac deposition in SMS-CTR cells. (E) Peak distribution of H3K27ac signals in SMS-CTR shScramble, SIX1 KD5, and SIX1 KD6 cells across promoters (+/-2.5kb from annotated TSS), 5/3’ UTR, gene body, and distal intergenic/enhancer regions. (F) ROSE analysis performed on shScramble and SIX1 KD H3K27ac ChIP peaks depicts the shift in *HEYL* (down) and *TNNT2/TNNI1* (up) SE rank between shScramble and differentiated SIX1 KD cells. Although not defined as an SE (top right quadrant of hockey stick plot), the LGR5 enhancer also shifts downward in SIX1 KD cells and is a gene associated with self-renewal and stem properties. (G) Pathway enrichment of genes associated with SEs identified in shScramble and the union of SEs identified in SIX1 KD5 and SIX1 KD6 (SIX1 KD) cells using gene set sizes restricted to 100 to 250 genes and a q-value cut-off of 0.05. (H) H3K27ac and SIX1 ChIP signal over the *HEYL* and *TNNT2/TNNI1* SEs depict changes in SIX1 binding abundancy at stem cell (*HEYL)* and muscle differentiation (*TNNT2/TNNI1)* loci during SIX1 KD-induced differentiation and respective levels of HEYL and TNNT2/TNNI1 expression as observed in the RNAseq data. ChIPseq tracks were generated using the Washington University Epigenome Browser.

### SIX1 globally regulates both stem/oncogenic and myogenic differentiation genes through fine- tuning of super-enhancer activity

To decipher the mechanism by which SIX1 loss results in transcriptional reprogramming, causing RMS cells to differentiate and stop growing, we performed an initial TF motif analysis using the RCisTarget R package to identify transcriptional regulators with predicted binding within +/-2.5kb of the TSS of the subset of differentially expressed genes. From this analysis, we observed strong enrichment for E-box motifs of which 41% (350/853) of the genes with expression differences with SIX1 KD were predicted to be regulated by the E-box myogenic TFs, *MYOD1* and/or *MYOG*, yet only 4% (37/853) of these genes were predicted to be directly regulated by SIX1 (Suppl Fig 6). Thus, we hypothesized that SIX1 loss leads to differentiation of RMS cells via reprogramming of myogenic TFs.

**Figure 6.**
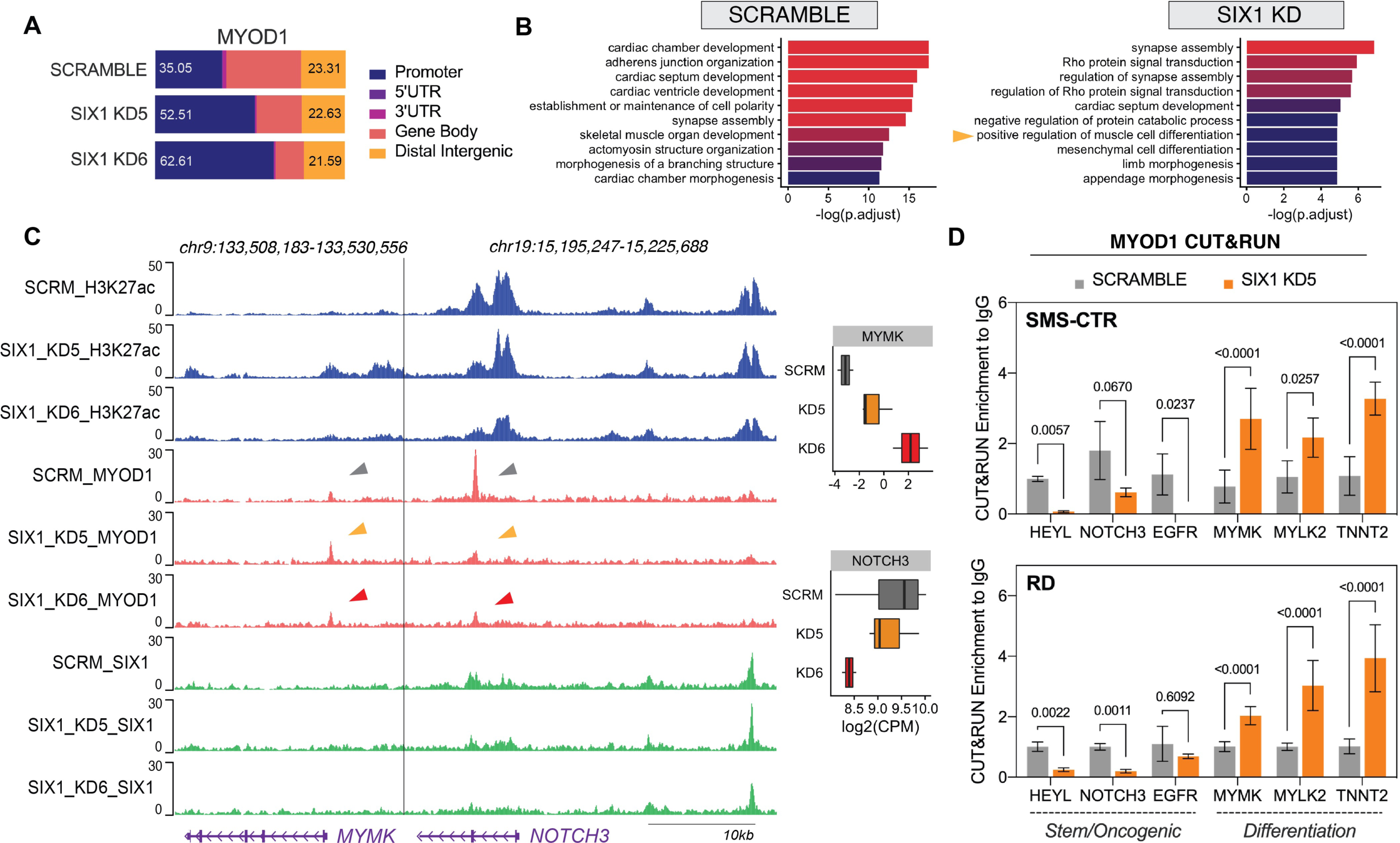
SIX1 loss alters MYOD1 occupancy at muscle differentiation and stem/oncogenic loci. (A) Peak distribution of the MYOD1 TF in SMS-CTR shScramble, SIX1 KD5, and SIX1 KD6 cells across promoters (+/-2.5kb from annotated TSSs), 5/3’ UTR, gene body, and distal intergenic/enhancer regions. (B) Pathway enrichment of annotated MYOD1 peaks that were called in shScramble and the union of MYOD1 peaks called in SIX1 KD5 and SIX1 KD6 cells (SIX1 KD). (C) H3K27ac, MYOD1, and SIX1 ChIPseq tracks over the *MYMK* and *NOTCH3* loci depict changes in MYOD1 binding that occur downstream of SIX1 loss and correlate with upregulation of MYMK and downregulation of NOTCH3 expression. ChIPseq tracks were generated using the Washington University Epigenome Browser. (D) CUT&RUN qPCR validation of changes in MYOD1 binding at stem/oncogenic (*HEYL, NOTCH3, EGFR)*, and myogenic differentiation genes (*MYMK, MYLK2, TNNT2)* that occur in SMS-CTR and RD SIX1 KD5 cells. Statistical differences were measured using a two-way ANOVA test followed by a *post- hoc* Sidak’s multiple comparisons test.

To determine how loss of the SIX1 TF activates a myogenic differentiation program, we performed chromatin immunoprecipitation followed by sequencing (ChIPseq) using a polyclonal antibody made against SIX1. We also performed ChIPseq against the master regulator of the myogenic lineage, MYOD1, and the active enhancer/chromatin histone mark H3-lysine-27 acetylation (H3K27ac) in SMS-CTR shScramble and SIX1 KD cell lines. Reflecting levels of shRNA-mediated SIX1 KD, we observed reduced genome-wide binding of SIX1 in both SIX1 KD lines compared to shScramble cells (Figure 5A) and sites of reduced SIX1 binding were highly enriched for SIX1/2 consensus motifs (Figure 5B). We further annotated genetic loci exhibiting 1.5-fold reduced SIX1 binding in both SIX1 KD lines compared to the shScramble control and found that SIX1 binding was reduced at gene loci involved in stem cell differentiation, Ras signaling, and cytoskeletal organization (Figure 5C). Accompanying sites of reduced SIX1 binding, we additionally observed decreases in MYOD1 and H3K27ac signal (Figure 5D, Suppl Fig 7). These data suggest that SIX1 predominantly plays a transcriptional activating role in FN-RMS and that SIX1 KD leads to a reduction in transcriptional output at stem-related and Ras-driven genes.

**Figure 7.**
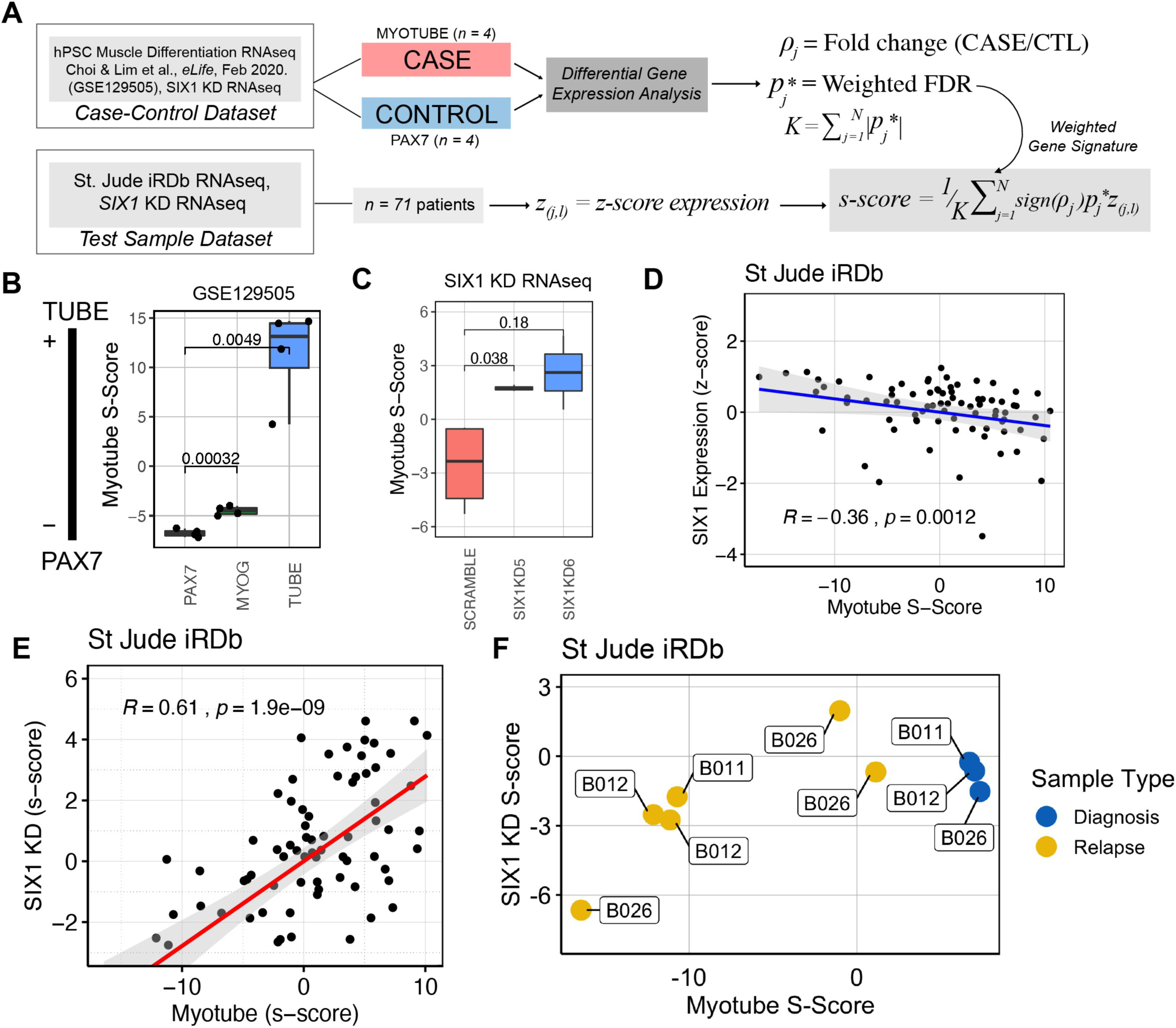
SIX1 expression in RMS is inversely correlated with a myotube gene signature. (A) Overview of *S*-scoring methodology whereby gene expression in the case-control (hPSC differentiated myotubes and Pax7+ progenitors, respectively) group is used to generate a weighted gene signature to score test sample transcriptomes on a continuous scale. (B) Myotube *S-*scores for samples used in training set plotted as proof-of-concept that the Myotube *S-*score can quantify myogenic differentiation status. Statistical differences measured by two-side Student’s *t-*tests. (C) Myotube *S*- scoring methodology applied to SIX1 KD RNAseq dataset demonstrates that SIX1 KD cells are more advanced in myogenic lineage than shScramble cells. Statistical differences measured by two-side Student’s *t-*tests. (D) Scatter plot of Myotube *S-*score plotted against SIX1 z-score-converted expression and Spearman rank correlation coefficient depict a moderate inverse correlation between differentiation status and SIX1 expression in St Jude iRDb RNAseq patient samples (*n =* 71). (E) Scatter plot of SIX1 KD *S-*scores derived from SIX1 KD RNAseq data against Myotube *S*-score shows a strong positive correlation between the SIX1 KD and myotube gene signatures in the St Jude iRDb expression dataset. (F) Myotube and SIX1 KD *S-*scores of three patient tumors (SJRHB011 = “B011”, SJRHB012 = “B012”, SJRHB026 = “B026”) collected and sequenced at multiple disease stages.

Given the changes in H3K27ac deposition and MYOD1 binding upon SIX1 KD, we hypothesized that SIX1 likely regulates large-scale transcriptional programs through mechanisms beyond direct transcriptional induction of *cis* genes. To determine how SIX1 KD affects global transcriptional output, we annotated H3K27ac signal distribution over promoters (+/- 2.5kb from TSS), gene bodies, and distal enhancers. In SIX1 KD cells, H3K27ac distribution increased at promoter regions (+/-2.5kb of TSS) and was reduced along gene bodies and moderately reduced at distal intergenic regions/enhancers (Figure 5E), showing a potential function of SIX1 in regulating enhancer activity in addition to promoter-based transcription.

To examine whether SIX1 levels influences enhancer activity, we compared enhancers and super- enhancers (SEs) via ranked H3K27ac signal between shScramble and differentiated SIX1 KD cells. Overall, 4.14%, 5.24%, and 7.37% of total H3K27ac peaks in shScramble (1470), SIX1 KD5 (1452), and SIX1 KD6 (1322) cells respectively corresponded to super-enhancers, which are characterized by long- ranging (over 12.5kb) clusters of strong H3K27ac deposition^29, 30^. Of note, we found that many oncogenic and myogenic genes marked as differentially expressed upon SIX1 KD in our RNAseq dataset were associated with SEs. For example, in SIX1 KD cells, we observed a downward shift in ranked H3K27ac signal at the SE associated with the Notch effector and muscle stem cell enriched gene *HEYL*^31^, and an upward shift of H3K27ac signal at the SE associated with the contractile muscle genes *TNNT2* and *TNNI1,* denoted as the *TNNT2* SE by the Rank Ordering of Super Enhancers (ROSE) algorithm (Figure 5F). We further annotated shScramble and SIX1 KD SEs by closest neighboring genes and discovered that although SEs occurred at myogenic-associated genes in both conditions, myogenic SEs in both SIX1 KD cell lines were associated with structural and contractile functions of skeletal muscle whereas those in the shScramble cell line were associated with less differentiated skeletal muscle pathways (Figure 5G). Side-by-side comparison of H3K27ac and SIX1 binding tracks at the example *HEYL* and *TNNT2/TNNI1* SEs not only reflects the shifts in SE activity seen in Figure 5F, but also demonstrates that SIX1 occupancy follows the pattern and trend of H3K27ac deposition (Figure 5H). Intriguingly, despite having global reduction in SIX1 binding and H3K27ac signal overall (Figure 5A, Suppl Fig 7), SIX1 binding and H3K27ac deposition at the *TNNT2/TNNI1* SE increased in the SIX1 KD5 line and remained relatively unchanged in the SIX1 KD6 line in comparison to that of the shScramble line. In contrast, SIX1 and H3K27ac signal at the *HEYL* SE were consistently reduced in both SIX1 KD lines, which contributed to the downward shift in *HEYL* SE rank in both SIX1 KD lines and reduced HEYL expression (Figure 5F&H). When examining the effects at a more global level, we observed similar reductions of H3K27ac signal at SEs associated with stem-related genes and relatively unchanged H3K27ac signal at SEs associated with muscle differentiation (Suppl Fig 7B). These data suggest that a loss of SE activity at stem genes may be the driving force of differentiation during SIX1 KD. As the SIX1 antibody used in ChIP has been shown to cross-react with other highly related SIX family members^32^, we reason that the lack of a decreased SIX1 binding at the *TNNT2/TNNI1* could be due to differences in SIX1 affinity to the myogenic loci during the differentiation state, or to the presence of a compensatory SIX member which could be recognized by the ChIP antibody. Nonetheless, these findings in the context of SIX1 loss of function, demonstrate a role for SIX1 in fine-tuning the activity of myogenic SEs that govern myogenic commitment as well as differentiation into contractile fibers.

### SIX1 loss alters MYOD1 occupancy at muscle differentiation and stem/oncogenic loci

By regulating SE activity, we reasoned that accessibility of myogenic TFs at oncogenic and myogenic loci could be affected by SIX1 KD. We observed that loss of SIX1 resulted in a change in MYOD1 distribution from distal intergenic/enhancer to promoter regions (Figure 6A). This coincides with the change in H3K27ac, indicating that SIX1 loss alters transcriptional dynamics, resulting in enhanced promoter-based and reduced enhancer-based transcription. Nearest gene annotation of MYOD1 peaks in the shScramble cell line and overlapping MYOD1 peaks in the SIX1 KD cell lines demonstrated that MYOD1 remains bound to myogenic loci in both shScramble and SIX1 KD genomes (Figure 6B). However, in the setting of reduced SIX1, we observed that MYOD1 sites occupied loci involved in positive regulation of muscle differentiation which did not appear in the top 10 pathways of shScramble MYOD1 peak (Figure 6B). Examples of the shift in MYOD1 binding are shown at the *MYMK* and *NOTCH3* loci (Figure 6C). Particularly at the *MYMK* locus, which is a key gene involved in myoblast fusion and formation of multinucleated myotubes^28, 33^, MYOD1 binding occurs at the gene promoter, and increases upon SIX1 KD (Figure 6C), which is consistent with its increased expression in SIX1 KD cells (Figure 4A&D, 6C). At the *NOTCH3* loci, MYOD1 binding occurs 22kb downstream of SIX1 within the *NOTCH3* promoter region and dramatically decreases upon SIX1 KD without a significant reduction in SIX1 binding at the upstream enhancer site and is coupled with downregulated mRNA expression (Figure 6C). In both these cases, we observed that changes in MYOD1 occupancy, rather than SIX1, aligned with H3K27ac marks. To validate the shift in MYOD1 occupancy at differentiation and progenitor-related genes in both SMS-CTR and RD cells, we performed MYOD1 Cleavage Under Targets and Release Under Nuclease (CUT&RUN) followed by qPCR (C&R qPCR), which is an orthogonal method to ChIP to detect target protein binding on DNA and requires far less cells than traditional ChIP methods^34^. We found that in differentiated SIX1 KD SMS-CTR cells as well as RD cells, MYOD1 was more abundantly bound at loci associated with differentiation genes, as opposed to myoblast or oncogenic genes (Figure 6D). These results reflect similar observations of MYOD1 genomic occupancy shifting as a consequence of myoblast formation or RMS induction toward differentiation^3, 13, 14, 35, 36^. Thus, our data demonstrate that SIX1 regulates a large-scale proliferative and less differentiated cell-identity program in RMS by maintaining MYOD1 binding at SEs resulting in a loss of promoter-driven myogenic gene transcription. Thus, SIX1 loss leads to an altered myogenic TF DNA binding landscape to one that is more permissive to the expression of contractile muscle genes over the expression of stem-related genes regulated by SEs.

### SIX1 expression is inversely correlated with a Myotube gene signature in RMS patients

The profound myogenic transcriptional program induced upon SIX1 inhibition suggests that the overexpression of SIX1 may serve as an upstream orchestrator of the aberrant muscle differentiation observed in RMS, as it does in normal muscle development^8^. To test this, we examined whether SIX1 expression in RMS patient samples correlates with an early myogenic transcriptional landscape. Using a recently published human pluripotent stem cell (hPSC) dataset^37^ aimed at defining the transcriptional landscape at multiple stages of human myogenic differentiation, we derived a myogenic differentiation signature from PAX7+ skeletal muscle progenitors and their final cell states as multinucleated myotubes. With this hPSC data to serve as case-controls for differentiated muscle and muscle progenitors, respectively, we applied a signature scoring method (*S*-score) previously described by Hsiao and colleagues^38^ to quantitatively score test data, RMS patient RNAseq samples, on their concordance to the gene expression signatures derived from empirical myotube-progenitor data (Figure 7A). To test the performance of our *S-*scoring methodology, we confirmed using the case-control hPSC data that *S-*scoring could segregate PAX7+ progenitors, MYOG+ myoblasts, and differentiated myotubes in a stepwise manner whereby the MYOG+ cells displayed an intermediate *S-*score between muscle progenitors and myotubes (Figure 7B). Furthermore, we calculated an *S-*score for our SIX1 KD RNAseq samples based on the myotube signature and were able to distinguish shScramble from SIX1 KD RMS cells based on this scoring method. SIX1 KD cells demonstrated greater alignment with the myotube signature, consistent with the results of other enrichment scoring methods used previously in Figure 4 (Figure 7C). Importantly, using this quantitative scoring technique, we are able to assess what stage of the myogenic differentiation cascade our RMS cells lie.

We next assessed how SIX1 expression correlates with myotube *S-*scores in RMS patient samples. In the St Jude iRDb cohort, we found a modest and statistically significant inverse correlation between SIX1 expression and myotube *S*-Scores (Figure 8D, Spearman correlation: R = -0.36, *p = 0.0012*). We additionally applied the same signature scoring algorithm to generate a SIX1 KD signature using our SIX1 KD RNAseq dataset as case (KD)-controls (shScramble) and *S-*scored both St Jude and GSE108022 RMS patient samples based upon SIX1 KD and myotube gene signatures. We observed strong positive correlations (St Jude: R = 0.57, *p<0.001*, GSE108022: R = 0.61, *p<0.001*) between the two signatures in the RMS patients, indicating that loss of SIX1 expression in RMS cells induces a transcriptional program highly similar to that which is observed by a myoblast transitioning to the myotube fate (Figure 7E).

Given the concordance of the SIX1 KD signature with the myotube signature, we next sought to examine whether these two signatures could be used to distinguish advanced RMS disease from primary disease. Of the 71 patient samples with complete RNAseq data available from the St Jude iRDb cohort, three of these patients had RNA-sequencing performed at multiple stages of the patient’s disease progression. Filtering down our analysis to these three patients, we examined whether disease recurrence was associated with changes in myogenic differentiation state. By myotube and SIX1 KD *S*-scoring, we observed that patient tumor expression profiles at diagnoses and disease recurrence states were distinguishable by differentiation and SIX1 KD scores, whereby relapsed tumors exhibited lower SIX1KD and myotube *S-*scores than their tumor at diagnosis (Figure 7F). Of note, we observed that the two relapsed tumor samples from patient B012 had lower Myotube and SIX1 KD *S-*scores compared to the tumor at diagnosis (Figure 7F). These data underscore our findings that the transcriptional program controlled by SIX1 in RMS is intimately linked to myogenic differentiation status, which is a driving force of RMS tumor progression.

## Discussion

Repression of myogenic differentiation programs is a known, critical attribute of RMS whereby dysfunctional MYOD1 and MYOG activity is thought to drive the disease^3, 39–42^. An unresolved question that persists in the field of RMS is why RMS tumors express the myogenic TFs, MYOD1 and MYOG, yet fail to progress past the apparent myoblast progenitor state^2, 43, 44^. While it is known that MYOD1 and MYOG have distinct subprograms that can drive either self-renewal or skeletal muscle differentiation, the departure of these MRFs from their canonical abilities to execute the complete sequence of skeletal muscle development in RMS invokes other factors that may repress the ability of MYOD1 to act on its differentiation programs. Therefore, the identification of other regulatory proteins that alter the context- specific functions of MYOD1 has become a core area of RMS studies^3–5^. Here, we report that the SIX1 homeobox TF acts as an upstream transcriptional regulator maintaining the arrest of RMS cells in a self- renewing muscle progenitor state. In the developmental context, the *SIX1* homeobox gene is highly expressed in early muscle development and is responsible for the direct activation of early MRF expression, but its expression becomes downregulated as the muscle reaches its final stages of differentiation^45, 46^. Using zebrafish and human cell line FN RMS models, we demonstrate that genetic inhibition of *SIX1/six1b* can trigger the activation of a muscle differentiation gene program in RMS cells, thus halting their growth and spread. These data are supported by preceding reports that show downregulation of *SIX1* occurs during the final stages of muscle differentiation and embryonic myogenesis^11, 37, 45, 46^, and further supports the hypothesis that aberrant SIX1 expression in RMS may be in part responsible for the MRF dysfunction occurring in RMS.

In the majority of studies implicating the role of *SIX1* in cancer progression, SIX1 ostensibly acts as a TF that induces the expression of downstream tumor-promoting genes. Notably, in two previous reports, the pro-metastatic functions of *Six1* in Rhabdomyosarcoma (RMS) were reported to be channeled through one of Six1’s transcriptional targets, Ezrin, a cytoskeletal protein^47, 48^, which was proposed to alter migration and invasion and thus contribute to RMS progression. In this study, we show for the first time that SIX1 promotes tumor growth/progression largely via alteration of global transcriptional programs of muscle cell-identity. Thus, while direct targets such as Ezrin likely contribute to its aggressive functions in RMS, the major function of SIX1 in RMS progression appears to be through changing cell fate by regulating transcriptional programs upstream of myogenic TFs. In normal development, *Six1* loss in muscle precursor cells leads to reduced MRF expression and concomitant defects in skeletal muscle formation^7, 8, 11, 16, 17, 49, 50^. In the context of FN-RMS, we observe that SIX1 KD is associated with loss of progenitor gene expression but a gain of muscle differentiation gene expression, raising the question of how SIX1 activates a differentiation program while it is itself suppressed. By ChIPseq, we observe that genome wide SIX1 binding closely overlaps with H3K27ac marks at promoters and SE regulatory elements. SIX1 KD leads to decreases in SIX1 binding at cytoskeletal, cell division, and stem-related loci, which aligns with previously characterized roles of SIX1^47, 48, 51–54^. On a global scale, SIX1 binding is enriched at SEs, enhancers, and promoters associated with cell division, cell-identity, and muscle specification. Upon SIX1 KD, SE activity as approximated by H3K27ac signal is diverted from progenitor/stem-related SEs to SEs associated with that of forming contractile muscle and other structural components of skeletal muscle differentiation, which manifest as the multinucleated and elongated morphology of SIX1 KD cells. In addition to these direct forms of transcriptional regulation either at target loci or at distal regulatory elements, we found that SIX1 can indirectly influence the DNA binding activity of MYOD1 and possibly other myogenic TFs by modifying the landscape of active chromatin and consequently TF binding accessibility at differentiation loci.

Pluripotency and cell type determination are controlled by the occupancy of master TFs and cell-type specific TFs at enhancer regions governing cell fate decisions^29, 55^. Within the repertoire of muscle-lineage enhancers, several TFs, which based on our studies include *SIX1*, have come to light as master TFs that initialize the myogenic lineage by sitting poised at myoblast enhancer elements and then become overactive in the context of RMS^35, 36, 39, 42^. Notably, these factors include the developmental TFs *SNAI1/2* and *TWIST2*, which similar to *SIX1* are found at stem and myogenic enhancer elements in RMS and are drivers of EMT, cell migration, and tissue repair^13, 35, 36^. Our focused study of *SIX1* compounds on growing evidence that the composition of TFs at muscle-specific enhancers controls the differentiation state of RMS cells, which raises multiple outstanding questions. First, this raises the question of what factors cause SIX1 to become overexpressed in FN-RMS tumors, particularly given the absence of *SIX1* amplification or any common perturbation of the locus. Whereas *SIX1* has been identified as target downstream of the *PAX3-FOXO1* fusion, the mechanism leading to SIX1 overexpression in FN-RMS is less understood^56^. Second, our findings raise the question of how diverse driver mutations associated with FN-RMS impinge on similar myogenic epigenetic/transcriptional programs in similar fashion to the *PAX3-FOXO1* fusion protein in FP-RMS^57–59^. Notably, genome-wide PAX3-FOXO1 fusion binding establishes SEs at myogenic genes and recruits the co-activator proteins p300, BRD4, and Mediator^59^, and similar functions may apply to TFs like *SIX1* in FN-RMS . Finally, the collection of these studies raises the question of whether RMS cells can be irreversibly reprogrammed to follow the proper cascade of myogenic differentiation through targeting master TF activity. Although there are still many barriers facing the viability of TFs as pharmacological targets, dissection of mechanisms that modulate specific TF activities can potentially reveal druggable nodes that control cell-type specific transcriptional programs. For example, the requirement of an EYA phosphatase co-factor interaction with SIX1 to strongly activate downstream target transcription represents one targetable node to SIX1 activity that our group is actively interrogating^60–63^. Thus, it will be of future interest to determine whether the EYA phosphatase plays a similar role together with SIX1 in trapping RMS cells in a progenitor-like state.

In summary, our studies demonstrate that the SIX1 TF prevents FN-RMS from undergoing the cascade of myogenic gene expression leading to differentiation via the regulation of transcriptional output at stem versus myogenic genes. We show that FN-RMS differentiates into non-proliferative myotube-like cells following SIX1 inhibition, and that the differentiation program is achieved by a shift in MYOD1 binding and enhanced transcriptional activity from genetic loci that foster cell growth to loci that specify and drive the myogenic lineage. Altogether, these findings define an epigenetic function of SIX1 in balancing the growth and differentiation properties intrinsic to the myogenic lineage and ultimately demonstrate SIX1 as suitable therapeutic target in RMS.

## STAR Methods

### Key Resources

| REAGENT or RESOURCE | SOURCE | IDENTIFIER |
| --- | --- | --- |
| **Antibodies** |  |  |
| Mouse monoclonal anti-SIX1 (1229, 992) | In-house purified antibody | N/A |
| Rabbit polyclonal anti-SIX1/2 | Atlas Antibodies | HPA0011893; AB_1079991 |
| Rabbit polyclonal anti-H3K27ac | Abcam | ab4729; AB_2118291 |
| Rabbit monoclonal anti-MYOD1 | Abcam | ab133627; AB_2890928 |
| Mouse monoclonal anti-myosin heavy chain | DSHB | MF-20; AB_2147781 |
| Mouse monoclonal anti-PAX7 | DSHB | PAX7; AB_528428 |
| Rabbit polyclonal anti-phosphohistone H3 (pSer10) | Sigma-Aldrich | H0412; AB_477043 |
| Rabbit polyclonal anti-cleaved caspase 3 | Cell Signaling Technology | 9661; AB_2341188 |
| Normal Rabbit IgG | Cell Signaling Technology | 2729; AB_1031062 |
| Mouse β-TUBULIN | Sigma-Aldrich | T4026; AB_477577 |
| Mouse β-ACTIN | Sigma-Aldrich | A5316; AB_476743 |
| Mouse β-ACTIN-HRP | Abcam | ab49900; AB_867494 |
| **Bacterial and virus strains** |  |  |
| Subcloning Efficiency DH5α competent cells | ThermoFisher | 18265017 |
| **Biological samples** |  |  |
| Rhabdomyosarcoma with striated muscle tumor array | Biomax | SO2082b |
| **Chemicals, peptides, and recombinant proteins** |  |  |
| Polybrene | Millipore | TR-1003 |
| Phenol-red solution | Sigma | P0290 |
| Puromycin Dihydrochloride | Research Products Int. | P33020 |
| pAG-MNase | EpiCypher | 15-1116 |
| Fugene Transfection Reagent | Promega | E2311 |
| Tricaine (MS-222) | Sigma Aldrich | A5040 |
| ECL Western blot substrate | Pierce | 32106 |
| Digitonin | Millipore Sigma | 30-041 |
| Spermidine | Sigma Aldrich | S0266 |
| **Critical commercial assays** |  |  |
| MycoAlert detection kit | Lonza | LT07-418 |
| Direct-zol RNA prep kit | Zymo Research | R2052 |
| iScript reverse transcription kit | Bio-Rad | 1708841 |
| SsoFast EvaGreen supermix | Bio-Rad | 1725205 |
| Verso cDNA synthesis kit | ThermoFisher | AB-1453A |
| Taqman gene expression master mix | Applied Biosystems | 4369542 |
| Nuclei EZ prep kit | Sigma Aldrich | NUC101 |
| Dynabeads Antibody Coupling kit | ThermoFisher | 14311D |
| Concavalin A beads | EpiCypher | 21-1401 |
| Universal Plus mRNA-Seq library prep kit | Nugen | 0508 |
| KAPA HyperPrep ChIP library kit | Roche | KK8502 |
| NEBNext II Ultra library prep kit | NEB | E7645, E7600S |
| DNA Clean and Concentrator kit | Zymo Research | D4033 |
| **Deposited data** |  |  |
| SIX1 KD RNAseq | This paper | N/A |
| SIX1 KD ChIPseq | This paper | N/A |
| Pan-Sarcoma and normal tissue expression | Downloaded from Oncogenomics database | https://fsabcl-pob01p.ncifcrf.gov/cgi-bin/JK |
| Pediatric Sarcoma expression | Downloaded from St. Jude PeCAN portal | https://pecan.stjude.cloud/proteinpaint/study/pan-target |
| Rhabdomyosarcoma patient RNAseq | Downloaded from St. Jude Integrated RMS Database | https://pecan.stjude.cloud/proteinpaint/study/RHB2018 |
| hPSC muscle differentiation RNAseq | <sup>37</sup> | GSE129505 |
| **Experimental models: cell lines** |  |  |
| HEK293T | ATCC | CVCL_0063 |
| Human: RH30 | Mark Hatley<sup>64</sup> | CVCL_0041 |
| Human: RH3 (RH28) | Mark Hatley | CVCL_L415 |
| Human: RH4 | Mark Hatley | CVCL_5916 |
| Human: RD | Mark Hatley | CVCL_1649 |
| Human: RH36 | Mark Hatley | CVCL_M599 |
| Human: RH2 | Mark Hatley | CVCL_A460 |
| Human: SMS-CTR | Mark Hatley | CVCL_A770 |
| Human: SMS-CTR stable shScramble | This paper | N/A |
| Human: SMS-CTR stable shSIX1 KD5 | This paper | N/A |
| Human: SMS-CTR stable shSIX1 KD6 | This paper | N/A |
| Human: RD stable shScramble | This paper | N/A |
| Human: RD stable shSIX1 KD5 | This paper | N/A |
| Human: RD stable shSIX1 KD6 | This paper | N/A |
| **Experimental models: organisms/strains** |  |  |
| Zebrafish: AB | ZIRC | ZL1 |
| Zebrafish: *six1b<sup>oz1</sup>* | Sharon Amacher<sup>11</sup> | N/A |
| Mouse: NOD/SCIDγ | CU AMC Breeding Core | N/A |
| **Oligonucleotides** |  |  |
| For SYBR cDNA primer sequences, see Supplemental Table 1.1 | This paper | N/A |
| For SYBR CUT&RUN primer sequences, see Supplemental Table 1.2 | This paper | N/A |
| For Taqman Primer/Probe sequences, see Supplemental Table 1.3 | This paper | N/A |
| **Recombinant DNA** |  |  |
| *rag2-KRASG12D* | David Langenau<sup>19</sup> | N/A |
| *rag2-eGFP* | David Langenau | N/A |
| *pLKO.1-shSIX1 KD5* (3'UTR) | Functional Genomics Core | TRCN0000015233 |
| *pLKO.1-shSIX1 KD6* (CDS) | Functional Genomics Core | TRCN0000015236 |
| *pLKO.1-shScramble* | Addgene | 1864 |
| **Software and algorithms** |  |  |
| FastQC | Babraham Bioinformatics | https://www.bioinformatics.babraham.ac.uk/projects/ |
| BBDuk | Joint Genome Institute | http://jgi.doe.gov/data-and-tools/bb-tools |
| STAR | <sup>65</sup> | http://code.google.com/p/rna-star/ |
| edgeR | <sup>66</sup> | https://bioconductor.org/packages/edgeR |
| clusterProfiler | <sup>67</sup> | https://bioconductor.org/packages/clusterProfiler |
| RCisTarget | 68 | https://bioconductor.org/packages/RcisTarget |
| Bowtie2 (v.2.3.4.3) | 69 | http://bowtie-bio.sourceforge.net/bowtie2/index.shtml |
| Samtools (v.1.11) | 70 | http://www.htslib.org/ |
| Picard | Broad Institute | http://broadinstitute.github.io/picard/ |
| Bedtools | 71 | https://github.com/arq5x/bedtools2/releases |
| MACS2 | 72 | https://pypi.org/project/MACS2/ |
| ChIPseeker | 73 | https://bioconductor.org/packages/ChIPseeker/ |
| ngs.plot | 74 | https://github.com/shenlab-sinai/ngsplot |
| deepTools | 75 | https://github.com/deeptools |
| HOMER (v.4.11) | 76 | http://homer.ucsd.edu/homer/ |
| Rank Ordering of Super Enhancers (ROSE) | 29,30 | http://younglab.wi.mit.edu/super_enhancer_code.html |
| R version 3.6.3 | The R project | www.r-project.org |
| Python version 3.8 | Python | https://www.python.org/ |
| FIJI | ImageJ | http://imagej.nih.gov/ij |
| Prism 9 | GraphPad | www.graphpad.com |
| IGV 2.8.0 | Broad Institute | https://software.broadinstitute.org/software/igv/2.8.x |
| Signature Scoring Algorithm (S-score) | 38 | In-house R scripts |
| **Other** |  |  |
| Nunc LabTek Chamber Slide System | ThermoFisher | 154526PK |

## Resource Availability

### Lead Contacts

Further information and requests for resources and reagents should be directed to and will be fulfilled by the lead contacts, Heide L. Ford and Kristin B. Artinger

### Materials availability

This study did not generate new unique reagents.

### Data and code availability

Scripts and code generated during this study are available upon request. RNAseq and ChIPseq datasets generated in this paper are deposited on GEO (GSE173155). Clinical datasets analyzed in this study are provided in the Key Resources table.

## Method Details

### Clinical RNAseq Datamining

Clinical sarcoma expression data was obtained from the NCI Oncogenomics database managed by Dr. Javed Khan at the NIH. Clinical RMS RNAseq expression data was downloaded from the St. Jude PeCAN portal and Integrated Rhabdomyosarcoma Database.

### Zebrafish line maintenance

Zebrafish lines used in this study were maintained in compliance with the University of Colorado Anschutz Medical Campus IACUC guidelines and policies. The *six1b^oz^*^1^ mutant line used in this study was a generous gift from Dr. Sharon Amacher’s lab and crossed as heterozygotes to generate wildtype, heterozygote, and mutant homozygote progeny. Fish were genotyped as described previously described^11^.

### Zebrafish ERMS Studies

Zebrafish ERMS tumors were established using previously described methods by the Langenau Lab. *rag2-kRASG12D* and *rag2-eGFP* plasmids were linearized with NotI and purified using the Zymo Clean and Concentrator kit. Linearized DNA was diluted to a stock concentration of 100ng/µL and injected with phenol-red dye into the single-cell stage of embryos for a final concentration of 5pg/embryo per *rag2* plasmid. Zebrafish tumor initiation events were recorded at 36 days post-injection and every week thereafter until 180 days. Tumor area was measured weekly using a Leica epifluorescent stereomicroscope along with body length to adjust for changes in basal growth of fish.

### Zebrafish *in situ* hybridization on zRMS tissue

Zebrafish tumor and normal muscle control tissues were fixed in 4% PFA for 2 hours at room temperature (RT), rinsed with PBS, and embedded in 1.5% agar/5% sucrose solution. Agar-sucrose tissue blocks were flash-frozen in liquid nitrogen and subsequently cryosectioned on a microtome. Frozen sections were defrosted for 1h at RT then incubated overnight at 70°C in six1b probe (provided by Vladimir Korzh, Institute of Medical and Cellular Biology, A*STAR, Proteos, Singapore) diluted 1µg/ml in hybridization buffer (1X SSC buffer, 50% formamide, 10% dextran sulfate, 1mg/ml yeast tRNA, 1X Denhardt’s). Sections were then washed 3x30min at 70°C (Wash: 1X Saline Sodium Citrate (SSC) buffer, 50% formamide, 0.1% Tween-20) followed by 3x10min at RT in MABT (1X maleic acid buffer, 20% Tween- 20), and incubated 2 hours in blocking solution (MABT, 20% sheep serum, 10% Boehringer Blocking Reagent). Sections were then incubated overnight at RT in 1:2000 anti-digoxigenin antibody diluted in blocking solution, washed 4x20min at RT in MABT, then 2x10min wash in AP staining buffer (100mM NaCl, 50mM MgCl2, 100mM Tris pH9.5, 0.1% Tween-20), and stained overnight at 37°C in 3.5µl/ml nitro- blue tetrazolium (NBT), 2.6µl/ml 5-bromo-4-chloro-3’-indolyphosphate (BCIP), 10% polyvinyl alcohol in AP staining buffer). Slides were rinsed 2X in PBS+0.1% Tween-20, 2X in ddH2O, dehydrated through ethanol solutions, cleared in xylene and coverslipped in Permount.

### Whole-mount zebrafish embryo *in situ* hybridization

Whole-mount RNA *in situ* hybridization in zebrafish embryos was performed as previously described^77^. DIG-conjugated antisense probes (gifts from Simon Hughes’ lab) were T7 or T3 transcribed for *pax3a, myod1,* and *myogenin* from pCS2+ backbone plasmids. *Post-hoc* genotyping of ISH-stained embryos was performed by incubating single embryos in 300mM NaCl overnight at 65°C to reverse crosslinks.

DNA was purified from each embryo by phenol-chloroform extraction and genotyped as described previously^11^.

### Cell Culture and Cell lines

FP-RMS and FN-RMS cell lines used in this study were a generous donation from Dr. Mark Hatley. Cell lines manipulated in this study (SMS-CTR and RD) were maintained at 37°C and 5% CO2 in Dulbecco’s Modified Eagle Medium (DMEM) supplemented with 10% FBS and 1% penicillin/streptomycin. Cell lines were tested for mycoplasma (Lonza MycoAlert) at least twice per year and only mycoplasma-negative cell lines were used in this study. All cell lines were STR authenticated by the University of Colorado Cancer Center Tissue Culture shared resource.

Stable SIX1 KD was achieved in SMS-CTR and RD cell lines by lentiviral transduction of two pLKO.1- derived shRNAs targeting the SIX1 CDS, subsequently denoted throughout the text as SIX1 KD5 and KD6. Control pLKO.1 shScramble cells were also transduced alongside SIX1 KD cells. pLKO.1 shRNA plasmids were transfected into HEK293T cells (293T) along with pMD2G and psPAX2 envelope and packaging plasmids. Viral particles were collected from 293T cells 48-hours post-transfection, passed through a 0.45µm filter syringe, and treated with 6-8µg of polybrene prior to infecting target cells. 24- hours post-viral infection, cells were selected with 2.0µg/mL (SMS-CTR) or 1.0µg/mL (RD) puromycin in 10% FBS/DMEM for 1 week and maintained in half the aforementioned puromycin dose for remaining experiments.

### IncuCyte Cell Growth Assay

RMS cell growth was measured on an IncuCyte Zoom (Essen Bioscience) Live-Cell Analysis platform. For cell growth, cells were plated at a concentration of 2500 cells/well in a 96-well plate and imaged every 12 hours with a 4X objective. Cell growth was measured by percent confluence and results presented in this study are normalized to percent confluence at time point zero (% Confluence to Baseline).

### qRT-PCR

Cells were harvested for RNA using the Zymo Direct-zol RNA isolation kit and cDNA was synthesized using the Bio-rad iScript reverse transcription kit following manufacturer’s instructions. Real-time qPCR was performed using Bio-rad ssoFast Evagreen supermix on a Biorad CFX96 qPCR instrument. SYBR primers used in this study are detailed in Supplementary Table S1.1.

Zebrafish tissues were snap-frozen in Trizol reagent, allowed to thaw, and homogenized using a plastic pestle. Homogenized tissue was then harvested for RNA using the Zymo Direct-zol kit. cDNA was synthesized using the ThermoFisher Verso cDNA Synthesis kit and qPCR reactions were performed using Taqman Gene Expression Master mix on an Applied Biosystems StepOnePlus instrument. Taqman probes used in this study are detailed in Supplemental Table S1.3.

### Western Blotting

Whole cell protein extracts were harvested by lysing cells in RIPA buffer treated with protease inhibitors and further lysed via sonification. 20-50µg of whole cell lysates were boiled with sample buffer and run through a 10% polyacrylamide gel. After PAGE gel electrophoresis, gels were transferred onto PVDF membranes, blocked in 5% Milk/TBST, and incubated with primary antibodies diluted in 5%BSA/TBST overnight at 4°C. Blots were incubated with HRP-conjugated secondary antibodies raised against primary antibody species at a 1:1000 dilution and chemiluminescence detected with Pierce ECL Western Blotting substrate. Chemiluminescence was imaged using an OdysseyFc imaging instrument. Between all antibody incubations, blots were washed with 1X TBST.

### Immunocytochemistry

Cells were plated on 4-well chamber slide and fixed in 4% PFA/PBS for 10 minutes and permeabilized in 0.1% TritonX-100/PBS (PBST) for 30 minutes. Chamber slides were next blocked with 15% goat serum/PBST for one hour and incubated in primary antibody solution overnight. The following day, chamber slides were incubated with appropriate fluorophore-conjugated secondary antibodies and mounted with Vectashield mounting medium with DAPI counterstain. All washes between incubation steps were performed with 1X PBS. Mounted slides were imaged on an Olympus BX51 fluorescence microscope. For phH3 and myHC stains, staining was quantified by dividing the number of positively stained cells by the total number of nuclei per field of view. Multinucleated events or fusion indices were quantified by counting the number of nuclei enclosed within a single positively stained myHC unit. For all immunocytochemistry stains, data is represented as image measurements taken over at least three independent experiments with two or more biological replicates per experiment, and two or more fields of view per biological replicate.

### Mouse Studies

All mouse studies were performed in 6-8 week old immunodeficient NOD/SCIDγ (NSG) of mixed genders. For mouse xenograft experiments, 2x10^5^ cells suspended in a 200µL 1:1 matrigel:1X PBS suspension were subcutaneously injected into either the left or right flank of the mouse, with each mouse receiving both a shScramble and SIX1 KD injection on one flank. Tumor growth was measured weekly for 12 weeks using calipers or until tumors surpassed a tumor volume of 1000 mm^3^ (1cm^3^). All animal studies were performed according to protocols approved by the University of Colorado Institutional Animal Care and Use Committee.

### Immunohistochemistry

For zRMS studies, tumor-burdened fish were euthanized in ice-water, fixed in 4%PFA overnight at 4°C, washed in PBS for 24 hours, decalcified in 20% EDTA pH 8.0 for 24 hours, dehydrated in 70% EtOH, and paraffin-embedded. Paraffin-embedded tissues were cut into 10-15µm thick sections and stained with H&E or further processed for antibody staining.

For mouse xenografts following dissection, mouse tumor tissue was fixed in 4% PFA overnight, washed in PBS for 24 hours, and dehydrated in 70% EtOH prior to paraffin-embedment. For all downstream IHC stains (zRMS, mouse xenograft, human tissue array), slides were de-paraffinized and retrieved in either pH6 (Six1, myHC) or pH9 (Pax7) Tris/EDTA buffer. Slides were then peroxidase blocked with 3% hydrogen peroxide (in methanol) for 10min, blocked in serum-free blocking reagent (DAKO) and incubated with primary antibodies for 1hr at room temperature. Appropriate species’ secondary antibodies were then incubated for 30min and developed with DAB stain for 10min and counterstained with hematoxylin for another 8min.

### RNA sequencing and Analysis

Total RNA was isolated from SMS-CTR cells using the Zymo Direct-zol RNA Miniprep Kit and RNA integrity confirmed using TapeStation analysis. shScramble and SIX1 KD SMS-CTR RNA samples were submitted as biological triplicates except for SIX1 KD6 which was submitted as biological duplicates on account of its marked proliferative defects. 100ng of total RNA per sample was used to construct PolyA- selected RNA libraries for RNAseq and sequenced using paired end reads with 150 cycles on an Illumina NovaSEQ 6000 instrument. Read QC was performed using fastqc and reads were trimmed with BBDuk to remove Illumina adapter sequences and the first 12 bases on the 5’ ends. Trimmed fastqc files were aligned to the hg38 human reference genome and aligned counts per gene were quantified using STAR^65^. Differential gene analysis was performed using the edgeR package^66^. Gene Set Enrichment Analysis (GSEA) was performed under default settings using the clusterProfiler R package gseaplot function^67^. Normalized counts (CPM) were converted to z-scores prior to plotting and heatmaps were created using the pheatmaps R package (https://CRAN.R-project.org/package=pheatmap).

### Chromatin Immunoprecipitation (ChIPseq)

Human cells along with spike-in Drosophila S2 cells at a 1:10 ratio with human cells were fixed in 1% formaldehyde diluted in growth media for an incubation time of 15 minutes. Crosslinking was quenched with the direct addition of 1M Tris pH 7.5 and shaking for 15 minutes. Cells were gently scraped off plates, pelleted by centrifugation, washed in cold PBS and centrifuged again. Cell pellets were snap frozen in liquid nitrogen and nuclei were extracted from cell pellets (Sigma Nuclei Isolation Kit #NUC-101). Chromatin was fragmented in sonication buffer (50mM HEPES pH 7.5, 140mM NaCl, 1mM EDTA, 1mM EGTA, 1% Triton-X, 0.1% Sodium deoxycholate, 0.1% SDS) supplemented with protease inhibitor cocktail using a Branson digital sonifier instrument at 4°C with the following settings: 7 cycles of 30s ON and 1m OFF sonification at 50% intensity. Chromatin lysates were incubated with 10μg antibody-bound Dynabeads (Dynabeads: Fisher Scientific #14-311-D; see supplemental materials for antibody information) overnight and subsequently washed in buffers of increasing stringency: 2X sonication buffer, 1X high salt sonication buffer (sonication buffer with 500mM NaCl), 1X LiCl buffer (20mM Tris pH 8.0, 1mM EDTA, 250mM LiCl, 0.5% NP-40, 0.5% sodium deoxycholate), and 1X TE pH 8.0. Immunocomplexes were eluted in 1% SDS/TE buffer and transferred to Lobind DNA tubes (Eppendorf #13-698-790) at 65°C for 30 minutes and crosslinks were reversed overnight by incubating samples at 65°C. RNA and protein were digested by the addition of RNase and Proteinase K, and DNA fragments were finally purified using phenol-chloroform. ChIPseq libraries were assembled using the KAPA HyperPrep ChIP library kit following manufacturer’s settings and were sequenced on an Illumina Nextseq500 machine.

### CUT&RUN

500,000 cells/sample were harvested by scraping and were resuspended and washed twice in wash buffer supplemented with protease inhibitor cocktail (20mM HEPES pH 7.5, 150mM NaCl, 0.5mM Spermidine). Cells were adsorbed onto activated Concavalin A beads for 10 minutes and then incubated with antibodies O/N at 4°C. After antibody incubation, unbound antibodies were washed away with cold Digitonin buffer (wash buffer + 0.01% Digitonin) and pAG-MNase was added to each sample to produce chromatin fragments under targets for 10 minutes at room temperature. Cells were then cooled to 0°C and incubated with ice cold 100mM CaCl2 for 2 hours at 4°C. MNase digestion was terminated with the addition of a master mix of STOP buffer (340mM NaCl, 20mM EDTA, 4mM EGTA, 50ug/mL RNaseA, 50μg/mL Glycogen) and 0.5ng/ul *E.coli* spike-in DNA and incubated for 10 minutes at 37°C. DNA was finally purified using a column purification kit and subsequently used for library assembly. Antibody concentrations: 1:100 for rabbit IgG and 1:50 for MYOD1. CUT&RUN libraries were assembled using the NEBNext II Ultra Library Prep kit) and dual-index primers following manufacturer protocols. Library size distribution was assessed by TapeStation and libraries were subsequently used for CUT&RUN qPCR.

### ChIPseq Analysis

The quality of the fastq files was accessed using FastQC (https://www.bioinformatics.babraham.ac.uk/projects/) and MultiQC^78^. Illumina adapters and low-quality reads were filtered out using BBDuk (http://jgi.doe.gov/data-and-tools/bb-tools). Bowtie2 (v.2.3.4.3) was used to align the sequencing reads to the hg38 reference human genome and to the dm6 drosophila reference genome^69^. Samtools (v.1.11) was used to select the mapped reads (samtools view -b - q 30) and sort the bam files^70^. PCR duplicates were removed using Picard MarkDuplicates tool (http://broadinstitute.github.io/picard/). The normalization ratio of each sample was calculated by dividing the total number of mapped reads mapping to the Drosophila genome of each sample by the total number of mapped reads mapping to the Drosophila genome of the sample with the lowest number of reads. Using the normalization ratio, random sub-sampling of the reads was performed using samtools view - hs. Bedtools genomecov was used to create bedgraph files from the bam files^71^. Peaks were called using MACS2 (v2.1.2) with default parameters for narrow peaks (--gsize hs --qvalue 0.01)^72^. Average profiles were generated using ngs.plot^74^ and heatmaps were generated using bigwig files with deepTools^75^. ChIP peaks were annotated using the ChIPseeker R package^73^. Super-enhancers were identified using the Ranking Ordering of Super-Enhancer (ROSE) algorithm using default parameters^29, 30^ and hockey stick plots were generated in R. ChIPseq track figures were generated using the Washington University Epigenome Browser^79^.

### Statistical Analysis

For all cell line experiments, experiments were performed in at least three independent biologi cal experiments with biological replicates and reported in this manuscript as a composite of these biological replicates. Therefore, when applicable, error bars for all figures including both cell line and animal experiments depict standard error of the mean (SEM). For all zebrafish experiments, an unpaired two- sided Student’s *t*-test was used to compare wildtype/control measurements to that of *six1b^-/-^* sibling or appropriately age-matched tumor tissue. For all cell line data, statistical differences between control and SIX1 KD conditions were measured using an unpaired two-sided Student’s *t-*test, unless specified otherwise in the figure legends. For animal experiments (both zebrafish and mouse) comparing tumor growth over time (Figure 2 & 3), tumor growth data were fitted to a Longitudinal Mixed Effect model and tumor growth was compared between shScramble and SIX1 KD mouse groups or wildtype and *six1b* mutant fish groups. Throughout this manuscript, all *p*-values are reported as is on figures or in figure legends.

## Supporting information

Supplemental Table 1: Primer sequences

Supplemental Table 2: GSEA output

## Acknowledgments

We would like to thank Dr. Brian Abraham for assistance with ChIPseq analysis and Erin Binne, Virginia Ware, and Taylor Hotz for their assistance throughout this project. We would also like to acknowledge a number of undergraduates whose help over the years contributed to this body of work: Julia Torline, Aaron Clark, Oscar Yip, and Hope Eden. This work was generously supported by NIH grants R21CA201809 (to H.L.F and K.B.A), R01CA224867 (to H.L.F), R01CA183874 (to P.J), K08CA245251 (to A.D.D), the Alex’s Lemonade Stand Foundation Innovation Award (to H.L.F), the CU Cancer Center Molecular and Cellular Oncology Pilot Grant P30CA046934 (to H.L.F, K.B.A, and P.J), and training fellowships T32GM763538 and TL1TR001081 (to J.Y.H). This work utilized the Cell Technologies, Functional Genomics, Pathology, and Biostatistics and Bioinformatics Shared Resource supported by P30CA046934 (to the University of Colorado Cancer Center). Research reported was also supported by the NIH/NCI P30CA021765 (to St. Jude Children’s Research Hospital Comprehensive Cancer Center), CureSearch for Children’s Cancer Foundation, Rally Foundation for Childhood Cancer Research, and the American Lebanese Syrian Associated Charities (to A.D.D and S.N).

## Diversity and Inclusion Statement

We worked to ensure sex balance/diversity in experimental samples through the selection of the cell lines, selection of non-human subjects, and selection of the genomic datasets (all which contained both male and female samples). The author list of this paper includes contributors from the location where the research was conducted who participated in the data collection, design, analysis, and/or interpretation of the work.

---

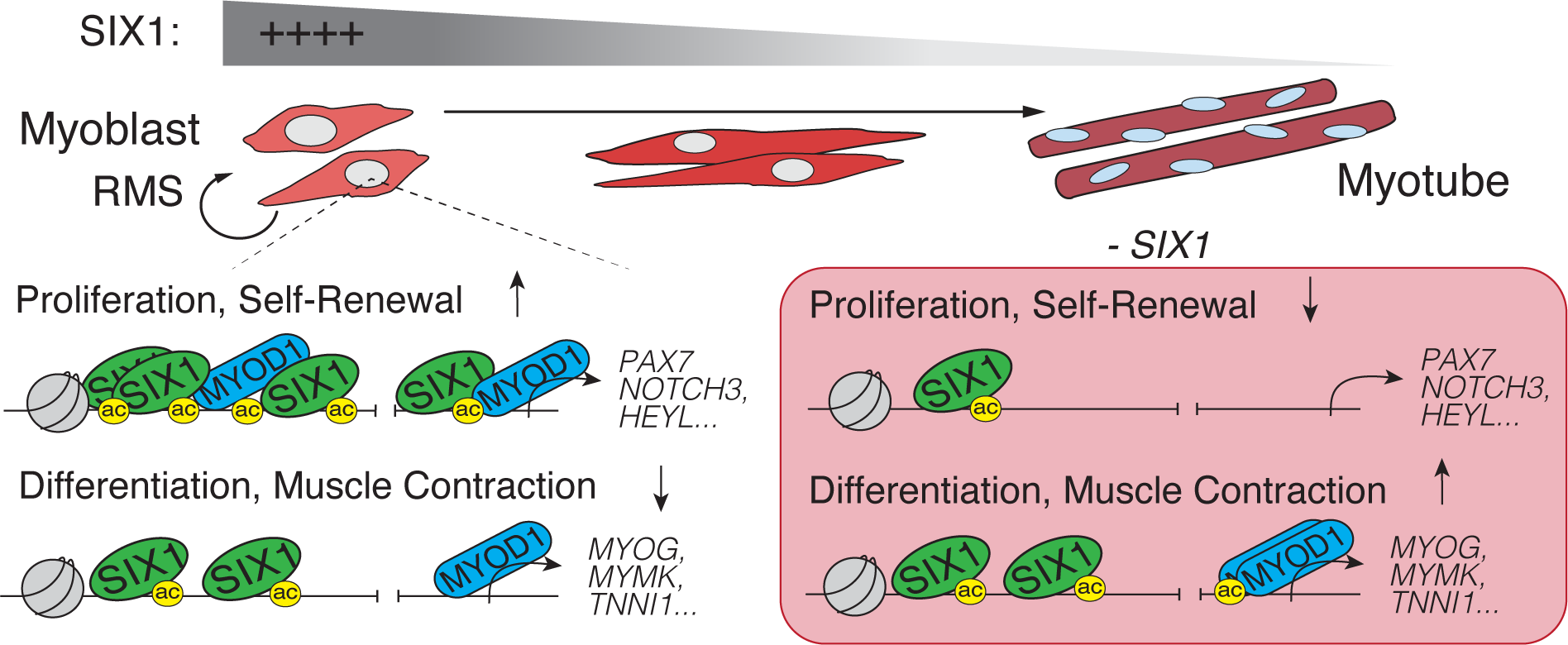

