## Supplemental Figure Legends for "SIX1 Reprograms Myogenic Transcription Factors to Maintain the Rhabdomyosarcoma undifferentiated state"

**Suppl Figure 1. SIX1 mRNA is highly expressed in human RMS**

(A) RMS (red) express higher levels of SIX1 compared to multiple sarcomas and normal tissue (grey) samples. Z-scored expression data was retrieved from the Oncogenomics database pan-sarcoma dataset. (SS = Synovial Sarcoma, RMS = Rhabdomyosarcoma, NBL = Neuroblastoma, OS = Osteosarcoma, MPNST = Malignant Peripheral Nerve Sheath Tumor, EWS = Ewing's Sarcoma, DSRCT = Desmoplastic Small Round Cell Tumor, ASPS = Alveolar Soft Part Sarcoma). (B) SIX1 TPM expression in RMS (red) and other pediatric sarcomas (black) from the St. Jude Pediatric Cancer Genome Project.

**Suppl Figure 2. *six1b* mutants have no overt defects in embryonic myogenesis**

(A) Western blot assessment of *six1* haploinsufficiency in *six1b*$^{-/-}$ adult skeletal muscle and age-matched wildtype skeletal muscle. Protein lysates were harvested from 1-year old wildtype and *six1b*$^{-/-}$ zebrafish. Data are a representative image of two independent experiments. (B-C) *In situ* hybridization (ISH) analysis of *pax3a, myod1,* and *myogenin* transcripts show *six1b* depletion does not affect expression in somite-matched wildtype and *six1b*$^{-/-}$ embryos across a time course of embryonic myogenesis. In-crossed embryos from *six1b*$^{+/-}$ breeding pairs were collected and subsequently probed for *pax3a, myod1,* or *myogenin* via ISH at the 5-7, 13-15 (*pax3a*), 8-10 (*myod1* and *myogenin*), and 20+ somite stages. Following visualization of all ISH stained embryos, genotyping was performed *post-hoc* to ISH staining to ensure blinded genotyping of the embryos during the ISH staining process (staining representative of $n \geq 5$ per somite stage and genotype).

**Suppl Figure 3. Reduced *in vivo* tumor growth upon SIX1 KD cannot be explained by differences in apoptosis**

(A) Representative images and quantification of cleaved-caspase-3 immunostaining (brown) counterstained with hematoxylin in SMS-CTR shScramble and SIX1 KD engrafted tumors. Each dot in graph represents %CC3+ staining per tumor section; 2 tumor sections quantified per tumor. Statistical analysis was performed using a two-sided Student's *t*-test.

**Suppl Figure 4. Positive and negatively enriched pathways upon SIX1 KD**

(A) GSEA plots of ranked log2FC RNAseq expression (SIX1 KD over shScramble) show negative enrichment of mesenchymal cell development and developmental cell growth gene signatures.

**Suppl Figure 5. Differentiated muscle histology in *six1b$^{-/-}$* zRMS tumor**

(A) Representative Pax7 staining (brown) and IHC quantification in wildtype and *six1b$^{-/-}$* zRMS tumor tissue (T) show overall weaker Pax7 staining in *six1b$^{-/-}$* tumor sections. Dots in graph represent %Pax7+ nuclei per tumor section; 3 tumor sections quantified per tumor-burdened fish. Statistical differences between wildtype and *six1b$^{-/-}$* tumors were calculated by a two-sided Student's *t*-test. (B) myHC PermaRed staining in wildtype and *six1b$^{-/-}$* zRMS tumor tissue show overall stronger myHC staining in one of 3 *six1b$^{-/-}$* tumor sections compared to weaker staining in wildtype tumor sections ($n$ = 4 wt tumors, $n$ = 3 *six1b$^{-/-}$* tumors).

**Suppl Figure 6. Transcriptional regulators identified from SIX1 KD differentially expressed genes**

(A) Top 10 results of the RCisTarget analysis performed on differentially expressed genes upon SIX1 KD. (B) Venn diagram of the 853 differentially expressed genes identified from the SIX1 KD RNAseq, 350 genes were predicted to be regulated by MYOD1 and/or MYOG and 20 of these genes were predicted to additionally be regulated by SIX1.

**Suppl Figure 7. Macs2 called peaks for SIX1, MYOD1, and H3K27ac in SMS-CTR shScramble and SIX1 KD cells and altered super enhancer activity at stem and myogenic SEs in SIX1 KD cells**

(A) Peaks called using default MACS2 narrowPeaks settings with q-value cut-offs of 0.01 for all ChIPseq data presented in this manuscript along with inputs. (B) H3K27ac (reads per million mapped reads) signal at super enhancers annotating to myogenic (GOBP: actin filament-based movement) and stem-related (GOBP: embryonic organ development) processes in shScramble and SIX1 KD cells.
