## Supplementary figures and images for "SIX1 Reprograms Myogenic Transcription Factors to Maintain the Rhabdomyosarcoma undifferentiated state"

### Supplemental Figure 1

#### Suppl Figure 1. SIX1 mRNA is highly expressed in human RMS tumors

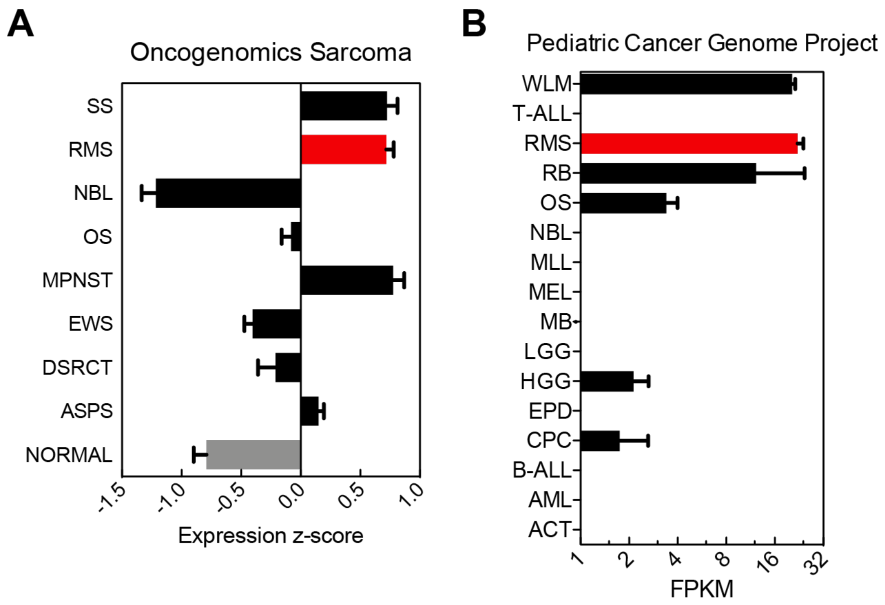

### Supplemental Figure 2

#### Suppl Fig 2. *six1b* homozygous mutants have no overt defects in embryonic myogenesis

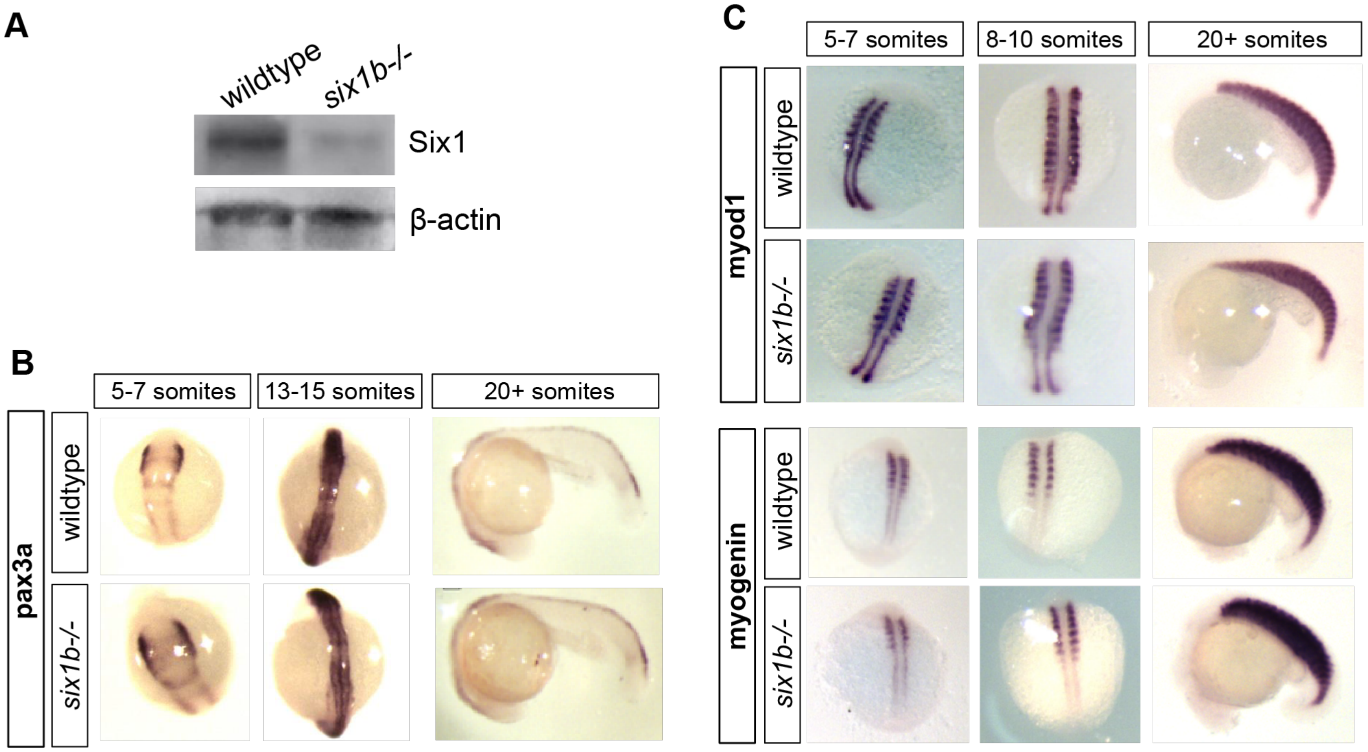

### Supplemental Figure 4

#### Suppl Fig 4. Positive and negatively enriched pathways upon SIX1 KD

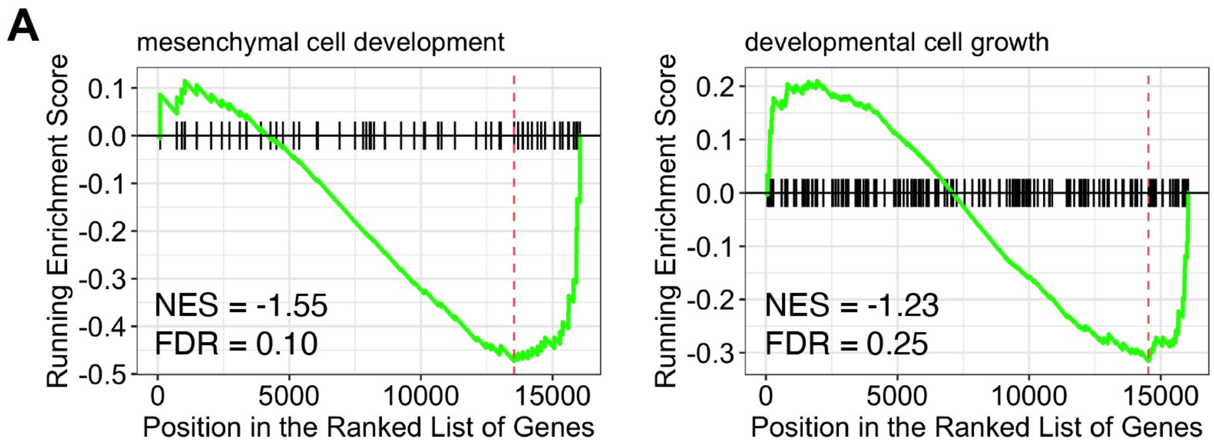

### Supplemental Figure 5

#### Suppl Fig 5. Differentiated muscle histology in *six1b-/-* zRMS tumors

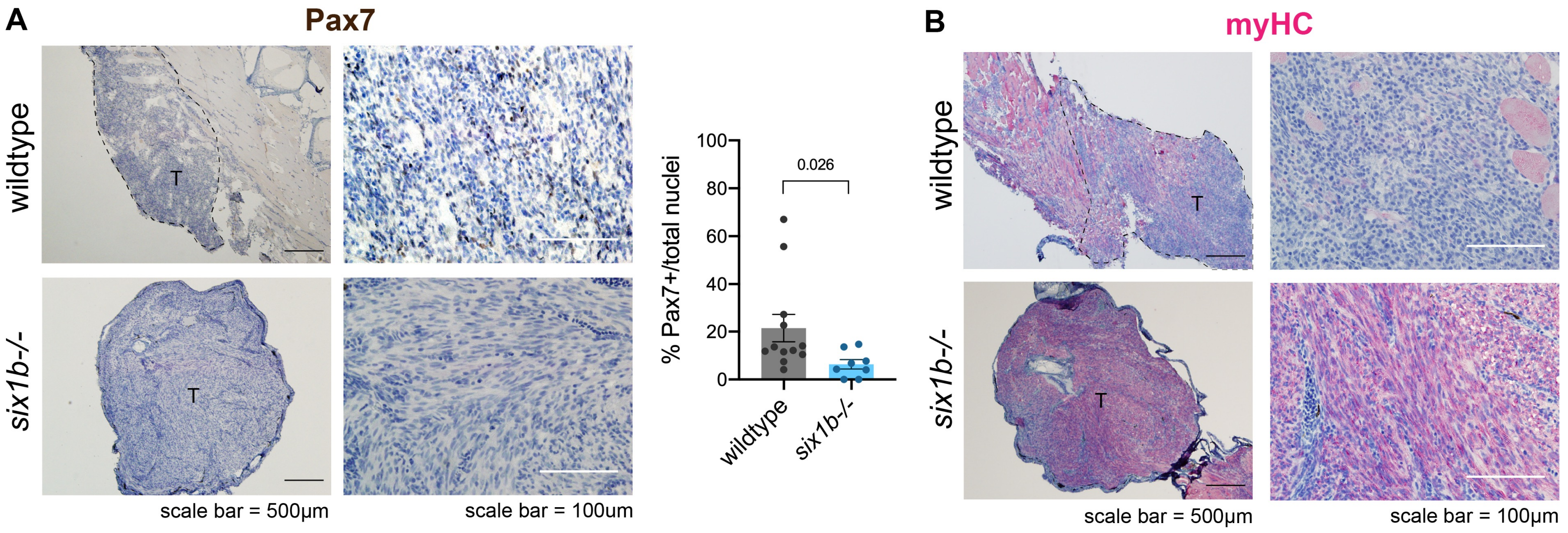
