## Supplemental Figure 3 for "SIX1 Reprograms Myogenic Transcription Factors to Maintain the Rhabdomyosarcoma undifferentiated state"

### Suppl Fig 3. Reduced *in vivo* tumor growth upon SIX1 KD cannot be explained by differences in apoptosis

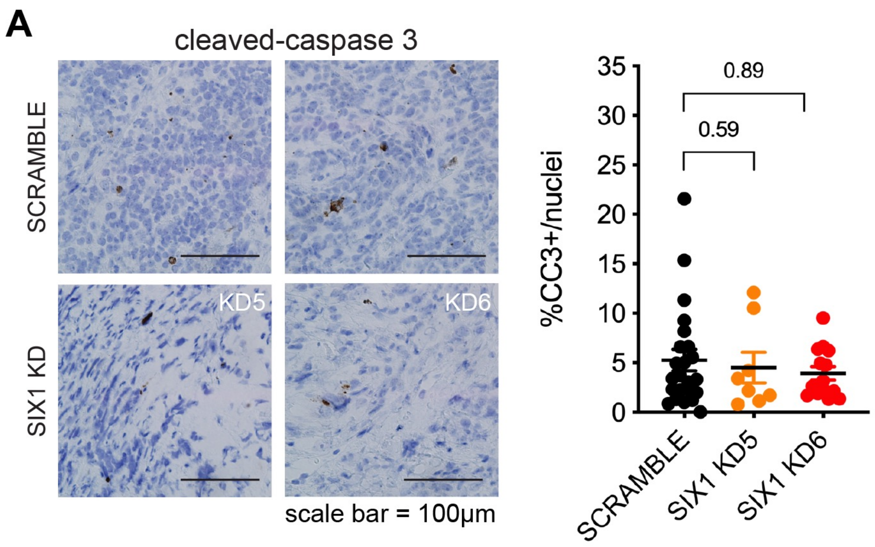
