## Supplemental Figure 6 for "SIX1 Reprograms Myogenic Transcription Factors to Maintain the Rhabdomyosarcoma undifferentiated state"

### Suppl Fig 6. Transcriptional Regulators identified from SIX1 KD differentially expressed genes

**A**

| Transcription Factor | Normalized Enrichment Score (NES) |
|---|---|
| MYOD1 | 6.61 |
| MYOG | 6.49 |
| MYF5 | 6.43 |
| TCF12 | 5.29 |
| ASCL2 | 4.96 |
| NEUROD1 | 4.79 |
| PITX1 | 4.79 |
| MYF6 | 4.66 |
| ASCL1 | 4.59 |
| TCF3 | 4.46 |

**B**

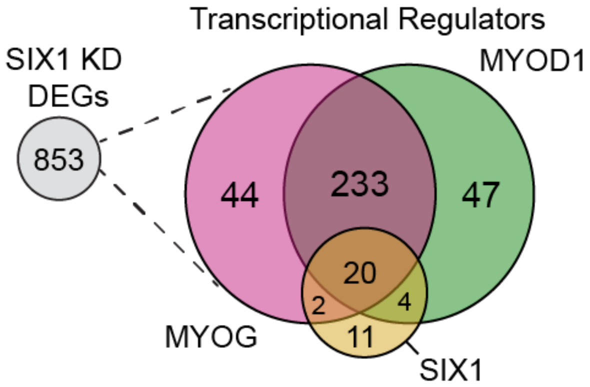
