## Supplemental Figure 7 for "SIX1 Reprograms Myogenic Transcription Factors to Maintain the Rhabdomyosarcoma undifferentiated state"

### Suppl Fig 7. Macs2 called peaks for SIX1, MYOD1, and H3K27ac in SMS-CTR cells and altered super-enhancer activity at stem and myogenic loci in SIX1 KDs

**A**

| Sample Name | Number of peaks |
|---|---|
| INPUT_SMS-CTR_SCRM | 443 |
| INPUT_SMS-CTR_SIX1KD5 | 68 |
| INPUT_SMS-CTR_SIX1KD6 | 84 |
| H3K27ac_SMS-CTR_SCRM | 35594 |
| H3K27ac_SMS-CTR_SIX1KD5 | 27812 |
| H3K27ac_SMS-CTR_SIX1KD6 | 18035 |
| MYOD1_SMS-CTR_SCRM | 3574 |
| MYOD1_SMS-CTR_SIX1KD5 | 686 |
| MYOD1_SMS-CTR_SIX1KD6 | 706 |
| SIX1_SMS-CTR_SCRM | 17723 |
| SIX1_SMS-CTR_SIX1KD5 | 19225 |
| SIX1_SMS-CTR_SIX1KD6 | 9673 |

**B**

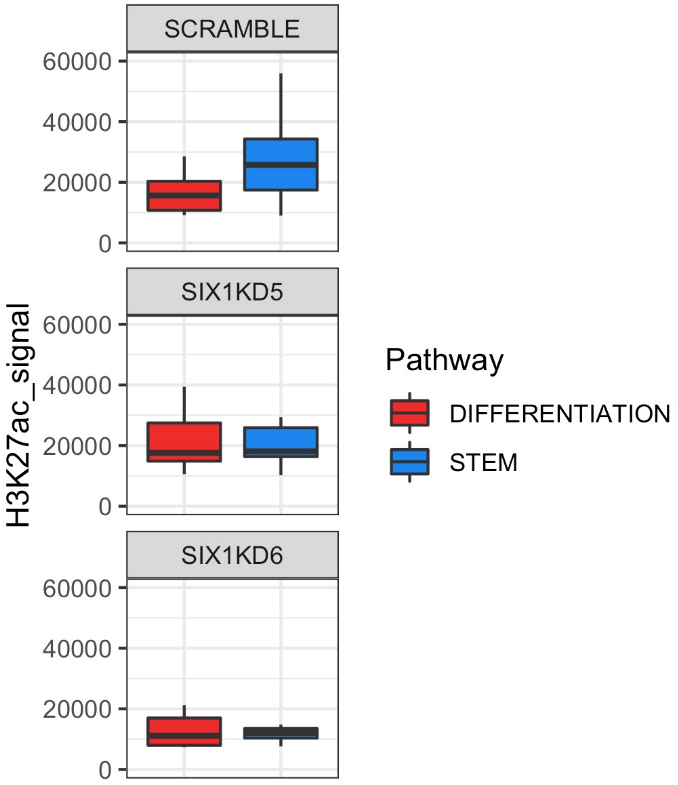
